# Associating EEG Functional Networks and the Effect of Sleep Deprivation as Measured Using Psychomotor Vigilance Tests

**DOI:** 10.1101/2024.05.22.595164

**Authors:** Sophie L. Mason, Leandro Junges, Wessel Woldman, Suzanne Ftouni, Clare Anderson, John R. Terry, Andrew P. Bagshaw

## Abstract

People are routinely forced to undertake cognitive challenges under the effect of sleep deprivation, due to professional and social obligations forcing them to ignore their circadian clock. However, low intra-individual and high inter-individual differences in behavioural outcomes are known to occur when people are sleep deprived, leading to the conclusion that trait-like differences to sleep deprivation could explain the differing levels of resilience. Within this study we consider if trait-like resilience to sleep deprivation, measured using psychomotor vigilance tests over a 40h constant routine protocol, could be associated with graph metrics (mean node strength, clustering coefficient, characteristic path length and stability) calculated from EEG functional networks acquired when participants are well rested (baseline). Furthermore, we investigated how stability (the consistency of a participant’s functional network over time measured using 2-D correlation) changed over the constant routine. We showed evidence of strong significant correlations between high mean node strength, low characteristic path length and high stability at baseline with a general resilience to extended sleep deprivation, although the same features lead to vulnerability during the period of natural sleep onset, highlighting non-uniform correlations over time. We also show significant differences in the levels of stability between resilient and vulnerable groups.

## Introduction

Sleep deprivation is endemic in society, for example the Centres for Disease Control estimate that in America more than one-third of adults are regularly sleeping insufficient amounts^1^. Partial and prolonged sleep restriction are both common forms of sleep deprivation, but while they differ in relation to sleep (i.e., some sleep vs. no sleep), for monophasic sleepers they share a common outcome - individuals are awake for longer periods of time, enhancing the homeostatic pressure for sleep. The effects of sleep deprivation, and associated fatigue, in humans are well documented both anecdotally and in studies. Sleep deprivation has been associated, in part, with major human and environmental disasters^2^. This includes the Chernobyl nuclear reactor meltdown, the ill-judged launch of the fatal Challenger space shuttle and the 2013 Metro-North train crash^3,4^. Within the US sleep deprivation was estimated to cost the economy $411 billion in 2015 due to higher levels of absenteeism (not attending work) or presenteeism (specifically attending work in a sub-optimal state for sleep related reasons)^5^. Sleep deprivation has a widespread impact on cognitive function, most typically captured through reaction times (RTs) on psychomotor vigilance tests (PVTs). General increases in RTs on PVTs as total time awake increases have been reproduced across many experimental protocols^6–9^. However, RTs are not simply a function of time awake but are controlled by the circadian rhythm of an individual, such as through the wake maintenance zone (WMZ) maximally promoting wakefulness before the release of melatonin^9^. Furthermore, compensatory effort can result in participants sporadically achieving performance equivalent to their baseline even after total sleep deprivation, known as the state instability hypothesis^8^.

However, large individual differences in the response to sleep deprivation are observed. For example, Van Dongen et al.^7^ and Chua et al.^6^ both repeatedly sleep deprived participants and observed large inter-individual differences in behavioural outcomes, whilst intra-individual results were relatively stable. Similar inter-individual differences in vulnerability were seen in active-duty fighter pilots on a flight simulator after 38 h of sleep deprivation, highlighting systematic differences even between highly trained professionals replicating a real-world task^10^. It was hypothesised by Van Dongen et al. that the different responses to sleep deprivation were due to individuals having a specific trait, which controlled their vulnerability to sleep deprivation^7^.

This has naturally led to studies trying to establish a biomarker for an individual’s vulnerability to sleep deprivation. For example, Van Dongen et al.^7^ considered both prior sleep history and baseline performance, but neither explained the trait-like vulnerability. However, Chua et al. found that individual differences in baseline measures (encompassing the first 16 h) in PVT response, blink rate, heart rate, heart rate variability and electroencephalogram (EEG) spectral power in the alpha band were each able to explain at least 50% of the variability between participants. This suggests that trait-like differences to sleep deprivation occur in electrocardiogram and EEG measures in addition to behavioural outcomes.

EEG holds a wealth of information regarding brain activity, and network properties could impact perturbations like sleep deprivation. Nonetheless, to the best of our knowledge, functional networks (FNs) derived from EEG data have not been used to assess susceptibility to sleep deprivation, despite neuroimaging being a potential biomarker source^11^. EEG combined with functional connectivity and network analysis is a powerful non-invasive tool for analysing the complex system of the brain at the macroscale and could therefore provide insights into how FN topology is associated with vulnerability to sleep deprivation. Indeed, it is plausible that the trait-like differences in the behavioural response to sleep deprivation could be underpinned by the topology of the individual’s FN, such that the robustness of this network to the perturbation of sleep deprivation drives inter-individual differences in behavioural outcomes. Therefore, in this study we use PVT performance from a 40 h constant routine (CR) to investigate if an individual’s response to sleep deprivation can be associated with information from their FNs obtained from well rested scalp EEG acquisitions before sleep deprivation. In particular, we consider the ability to correlate (i) an individual’s impairment (a singular metric of sleep deprivation effects) and (ii) an individual’s performance (continuous metrics across the CR) on a PVT, with graph metrics derived from FNs calculated using a baseline EEG acquisition. Furthermore, we use a stability analysis (the consistency of a participant’s FN over time measured using 2-D correlation) to further understand why PVT impairment differs between individuals.

## Methods

### Ethical Approval

Ethical approval was provided by the Monash University Human Research Ethics Committee (CF14/2790 – 2014001546). Participants were required to give informed consent before involvement in the study. All provided data were already anonymised; therefore, no personal information could be associated with an individual.

### Participants and Protocol

Exclusion criteria were (i) no artificial disruption to normal sleep patterns prior to the study (no shift work defined as 5 or more hours worked between 20: 00 and 07: 00 or transmeridian travel across three or more timezones) for at least the previous three months (ii) no prior diagnosis of medical, psychiatric or sleep disorders (iii) no extreme chronotype - the phase relationship between an individual’s circadian rhythm and the light-dark cycle of the sun. These criteria ensured participants were not experiencing sleep disruption or circadian misalignment prior to the CR. Participants were good healthy sleepers, reporting habitual bedtimes between 21: 30 and 01: 00 and habitual wake times between 05: 30 and 09: 00. This resulted in the study having 13 participants (*n*=2, female) 24.85 ± 4.26 years (mean ± standard deviation) where PVT and EEG data were available. EEG data for a further 9 participants was available. Baseline graph metrics were not dissimilar to those in the study (*p* > 0.05, t-test) and the stability across the CR for these participants is shown in the Supplementary Figure S22.

The study consisted of a six-day in lab protocol. After two days of acclimatisation within the laboratory (B1 and B2), participants awoke on the third day to a 40 h CR (CR1, CR2) followed by two days of recovery (12 h opportunity) sleep (R1, R2), as seen in Fig. 1. The CR consisted of 40 h continuous wake in private, sound-attenuated, temperature and light controlled suites. Participants were closely monitored to ensure sustained wakefulness and all time-cues were removed. Light (∼ 3 ± 1 lux (horizontal) and ∼ 1 ± 3 lux (vertical)) and temperature levels (21°*C* ± 2°*C*) were kept constant. In addition, participants remained in a semi-recumbent position throughout. Finally, isocaloric snacks were provided hourly, following Australian Dietary Guidelines 2013 for macronutrients^12^.

**Figure 1.**
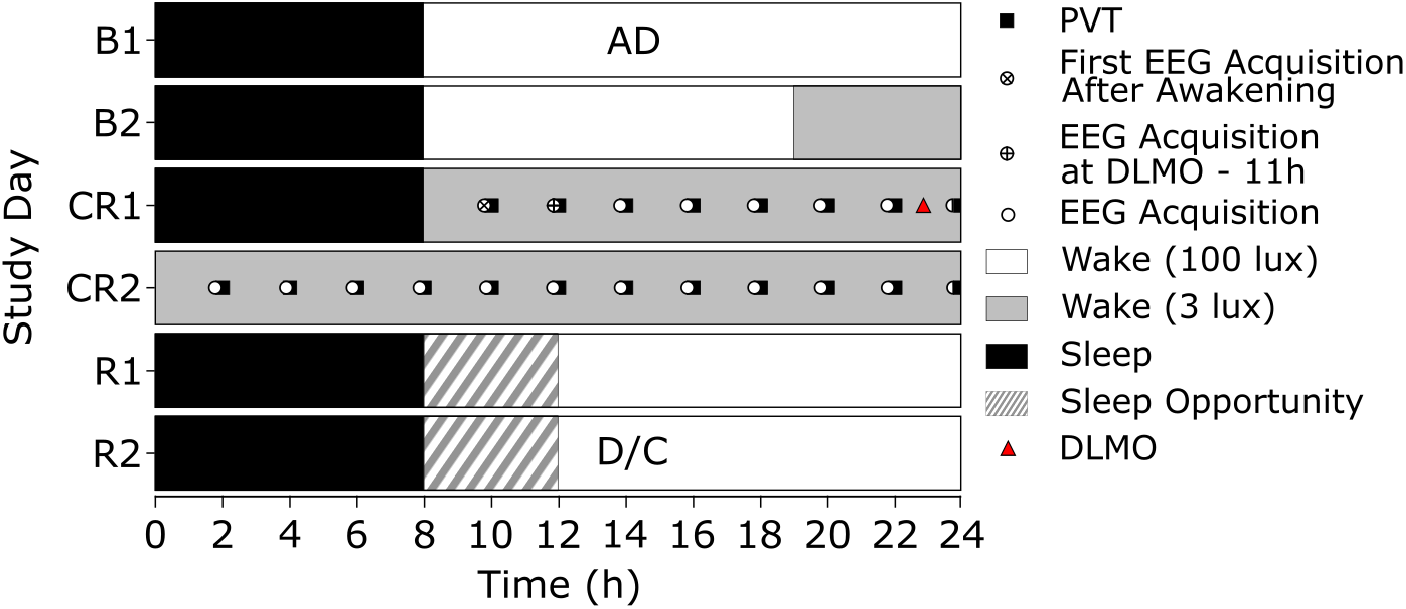
CR Protocol. Raster plot depicting the laboratory protocol across the two baseline days (B1, B2), the two days of CR (CR1, CR2) and two recovery days (R1, R2) - nominal 08:00 waketime and 23:00 DLMO as denoted by the red triangle. AD, admissions; D/C, discharge; circle EEG acquisition (150 s); circle and cross, first EEG acquisition after awakening (150 s); circle and plus, EEG acquisition at DLMO − 11 h (150 s) - here DLMO was set at 23:00, as denoted by the red triangle, to provide an example of the first acquisition after awakening differing from the EEG acquisition at DLMO − 11 h.

**Figure 2.**
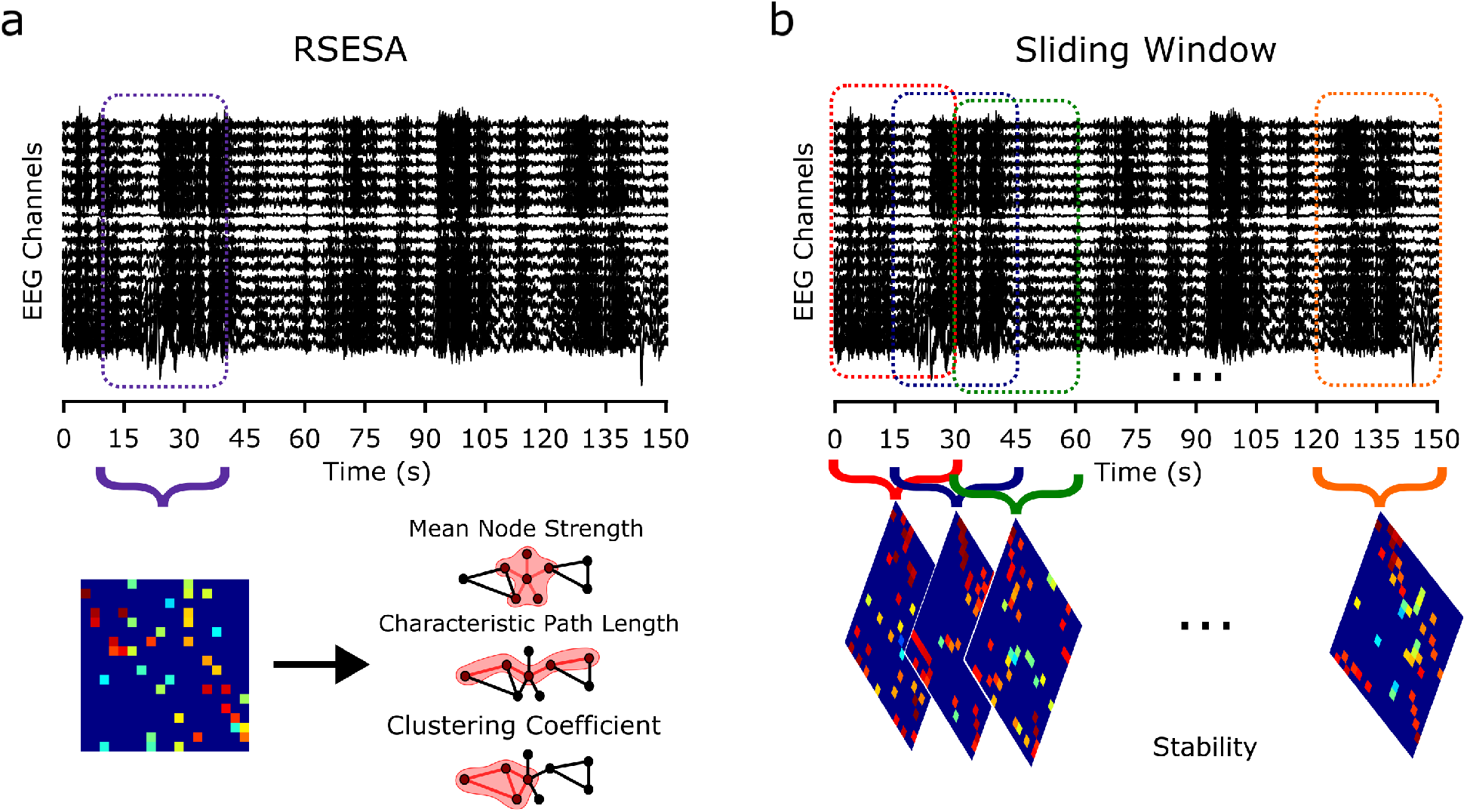
Epoch Selection. **(a)** A single 30 s epoch selected using the Resting State Epoch Selector Algorithm, which was used for calculating a FN and the graph metrics mean node strength, characteristic path length and clustering coefficient. **(b)** The sliding window approach used to select 30 s epochs with a 15 s overlap. For each epoch a FN is calculated and the median 2-D correlation between each pair of FNs was taken as representative of stability.

The experimental protocol ensured participants refrained from consuming alcohol and caffeine or using nicotine, supplements or prescription or non-prescription drugs for 3 weeks prior to the protocol beginning. This included women not using hormonal contraception or being pregnant. These measures were verified through a urine toxicology screen and a breathalyser test. In addition, for the three weeks prior to laboratory settings and the first two days in the clinic a strict 8:16 hours sleep:wake schedule was adhered to^13^. Compliance was checked via actigraphy data collected from Actiwatch Spectrum (Philips Repironics, BMedical, QLD, Australia) and time-stamped call-in messages upon waking and when participants went to bed. For a detailed description of the CR see McMahon et al.^9^.

### Dim Light Melatonin Onset Calculation

Melatonin levels were measured during the protocol for the calculation of Dim Light Melatonin Onset (DLMO) (21: 13±00: 58) to provide a marker of circadian phase per individual. Plasma melatonin levels for each participant were assessed through hourly blood samples starting approximately 1 h after starting CR. Values for hourly melatonin levels were then interpolated with DLMO times calculated as the first time that plasma melatonin concentration exceeded 5 pg/mL. Where possible DLMO was calculated for both the first and second evening of the CR. DLMO was only established using melatonin concentrations from the second evening when a recording for the first evening was not possible, for those with both nights the two nights were highly correlated *r* = 0.94. The circadian phase for each participant was then set, such that DLMO on the first day was defined as time 0. Participant PVT data and EEG data were aligned by selecting the acquisition closest to *n* hours before or after that participant’s DLMO. For example, when aligned by DLMO the EEG acquisitions at DLMO − 11 h took place −11: 05 ± 00: 39 hours before DLMO while the PVT took place −11: 17 ± 00: 31 hours before DLMO.

### Psychomotor Vigilance Test

Every two hours each participant undertook a 10-minute PVT. Here participants watched a white rectangle on the centre of the screen and had to respond with a thumb press as quickly as possible when the counter appeared. The stimulus for each trial appeared at a random interval of 2 to 10 s after the previous trial.

A RT under 100 ms was considered to be an anticipatory error and was removed from analysis in line with Basner and Dinges^14^. Trials with RT over 500 ms were labelled as lapses and were included in the analysis^14^. A RT over 5, 000 ms produced an alerting noise^9^ before the next trial started. The mean, median and std of RT, and the percentage of lapses from useable trials for each 10-minute test, were calculated.

The results of certain testing sessions were not used in the analysis due to the test either not occurring or lasting less than the required 10 minutes (*n* = 13 tests (5.26%) from *n* = 4 participants). The available tests for each participant are given in Fig. S1.

### PVT Impairment and Performance

Performance was monitored throughout the CR using the PVT, where the mean, median and standard deviation of RT and percentage of lapses were recorded (Fig. 3 and Supplementary Fig. S3 display these time-series). Hence, for each participant we have up to 19 datapoints across the CR for PVT performance in a given metric.

**Figure 3.**
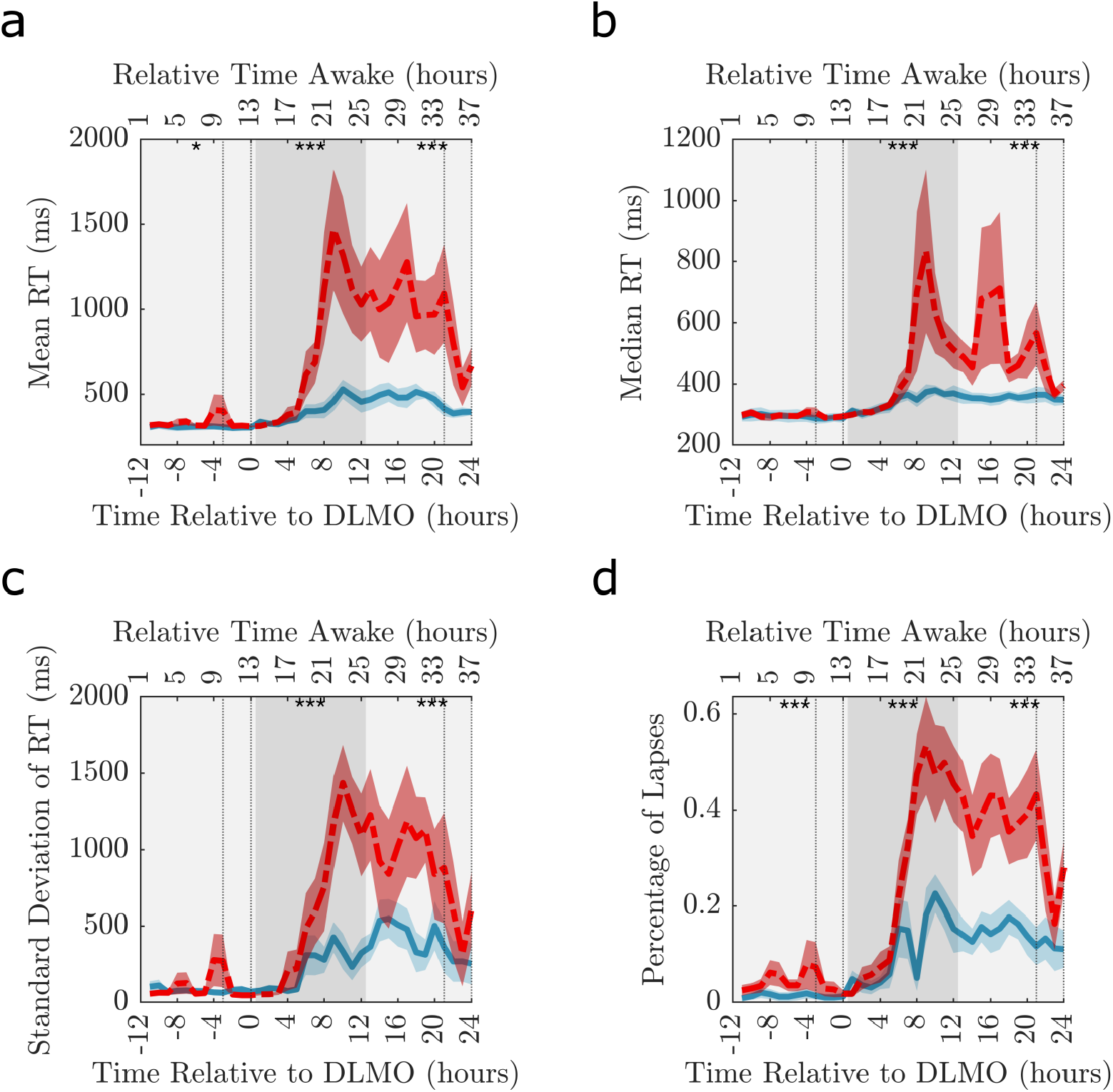
Psychomotor Vigilance Test Performance Split into Groups Based on High and Low Impairment. Timeseries showing **(a)** mean RT, **(b)** median RT, **(c)** standard deivation of RT and **(d)** the percentage of lapses when the participants are split into two groups based on the level of impairment in that measure. The solid blue line 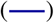 and dashed red line 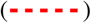 are the mean across the 6 lowest and 6 highest impaired participants, respectively. The corresponding blue and red shaded area is the standard error of the mean. The dotted vertical lines indicate the times considered to be in the WMZ (3 hours before DLMO to 5 minutes after). The significance of permutation tests (*n* = 10, 000) comparing the mean value of the low and high impairment groups over adjacent 12-hour periods depicted by different shades of grey are represented by, * *p* < 0.05, ** *p* < 0.005, *** *p* < 0.0001. In all plots the relative time awake provides an indication of the time awake for the majority of the participants.

We also wanted to detect impairment irrespective of when the worst performance occurred, removing inter-individual differences related to timing. For the mean, median and std of RT we calculated the magnitude of impairment as the percentage difference from the best performance prior to and include DLMO in CR1 (e.g., the biological day) to the worst performance post-DLMO (e.g., after the onset of sleep deprivation). For the percentage of lapses impairment was represented by their worst percentage of lapses post-DLMO. Hence, for each participant we have 1 value of PVT impairment in a given metric.

These time-periods were selected to separate the biological day from the onset of the biological night post-DLMO and hence the start of the extended sleep deprivation protocol. The timings of the PVT used for calculating impairment are presented in Supplementary Table S1 demonstrating that the timing of the best and worst performance can differ across both PVT metrics and participants.

The PVT impairment of each participant was then used to divide the participants into two groups. Hence, the 13 participants were split into the 6 lowest and 6 highest impaired participants for a given PVT metric, with 1 participant not included. The median participant was removed to allow greater differentiation between the high and low impairment groups and ensure equal group sizes. Note that the participants were split by the level of impairment for each measure, hence the high and low impairment group changed per PVT metric.

### EEG Acquisition

EEG data were recorded using 18 scalp electrodes from the standard 10-10 system (Fp1, Fp2, F3, F4, Fz, C3, C4, Cz, P3, P4, Pz, P7, P8, Po3, Po4, O1, O2, Oz). All EEG acquisitions were collected on a Compumedics Grael system with Profusion Sleep 4 software (Compumedics Ltd, Vic, Aus) with a sampling rate 512 Hz. The first 150 s recording occurred 2 h after waking and subsequent recordings occurred every 2 h for a total of 40 h awake. EEG data were collected under Karolinska Drowsiness Test (KDT) conditions^15^. Participants were instructed to keep their eyes closed and remained relaxed and still. KDT conditions ensured a clear segment of wake EEG with minimal blink/movement artifact. This resulted in up to 19 acquisitions per participant.

For acquisitions where one or more electrode time-series were corrupted, the entire acquisition was removed (*n* = 5 acquisitions (2.02%) from *n* = 3 participants covering *n* = 3 electrodes). This was to ensure all EEG networks contained the same base network of 18 nodes making each FN directly comparable. In addition, *n* = 6 (2.43%) EEG acquisitions from *n* = 3 participants were missing, due to technical issues with the recording devices or participants requiring medical attention or experiencing distress. Supplementary Fig. S2 shows the available EEG recordings for each participant.

### EEG Preprocessing

For each 150 s recording the EEG was referenced to the common average. The EEG acquisition was bandstop filtered between 49-51 Hz to remove powerline interference and filtered into the broadband (0.5-30 Hz). For the calculation of baseline graph metrics each epoch was filtered into the 8 − 14 Hz (alpha) frequency band as resting state eyes closed EEG has high alpha spectral power when the participant is well rested^16^ and because Chua et al.^6^ found that only the alpha frequency band significantly explained intra-individual variance in task performance after sleep deprivation.

However, for the stability analysis four frequency bands were considered, 0.5 − 4 Hz (delta), 4 − 8 Hz (theta), 8 − 14 Hz (alpha) and 14 − 30 Hz (beta), chosen to ensure each frequency between 0.5 − 30 Hz was included. Stability was considered across all four frequency bands as the measure was calculated across the CR and the power spectral density of each frequency band is known to change with time awake^17^. Within each frequency band the data were then normalised such that each channel had mean 0 and standard deviation 1^18^.

### Baseline EEG Acquisition

The baseline EEG acquisition was defined as either: B_*FA*_ the first EEG acquisition after awakening (two hours post wake) or B_−11_ the acquisition 11 hours before DLMO (between 2 and 4 hours post wake). These two baselines differ slightly by allowing an alignment based on time awake and internal phase, respectively. For the majority of participants B_−11_ was the same as B_*FA*_, however for 3 participants their EEG acquisition 11 hours before DLMO was acquired 4 hours post wake.

### Functional Network Construction

For the baseline acquisition 30 s epochs were utilised and the 18 channels of filtered and normalised EEG data were used to calculate FNs. The phase locking factor (PLF) was used to calculate the functional connectivity between pairs of nodes^19^. Each frequency band filtered EEG timeseries was considered an oscillator, with the position of oscillator *i* at time *t* ∈ {1,…, *L*} given by 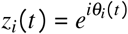, where *θ*_*i*_(*t*) is the oscillator’s phase. Then the level of synchronisation between pairs of nodes (e.g., nodes *j* and *k*) was summarised by how similar the phase of the two oscillators were,

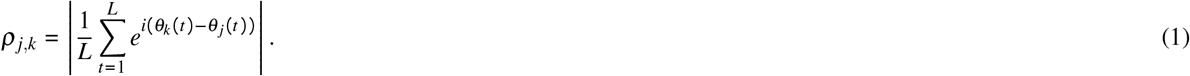

This resulted in undirected weighted adjacency matrices *ρ* ∈ ℝ, each with a size 18 × 18 and edge weights in the range [0, 1]. Surrogate analysis was then used to calculate the significance of each connection. 100 surrogate time-series were created by altering the bandpass filtered time-series using IAAFT - a Fourier-based method which ensures the autocorrelation and spectrum of the time-series were preserved^20^. The same methodology was used to create 100 surrogate PLF networks. Edges in the participant’s original PLF network remained if their weight was significantly higher than the corresponding edge from the surrogate networks at a 95% significance level, whilst non-significant edges were replaced by 0^21^.

In addition, the lag between each pair of EEG times series was calculated. Edges with a corresponding lag of zero were removed due to volume conduction^22^. Furthermore, the lag was used to make the surrogate-corrected PLF network directed by retaining edges with a positive lag^23^.

Finally, an order two edge reduction was used to remove connections where an alternative path via one or two additional nodes was stronger^21^. This process for creating FNs was chosen to reduce the impact of indirect connections.

### Calculating Graph Metrics

Four graph metrics were then calculated for each alpha baseline PLF network: mean node strength, clustering coefficient, characteristic path length and stability.

The mean node strength, clustering coefficient and characteristic path length analysis were calculated using Brain Connectivity Toolbox^24^ for one 30 s epoch per baseline acquisition. Hence, a single 30 s epoch was selected from each baseline 150 s acquisition (Fig. 2a). EEG acquisitions were collected under supervision within a strict KDT protocol^15^. As a result, the artifacts were minimal and multiple 30 s epochs could have been selected. To provide an objective way to select an epoch a Resting State Epoch Selector Algorithm (RSESA) was used. The algorithm choose an epoch by finding 30 s of the acquisition which maximised the peak alpha frequency in the occipital electrodes O1 and O2, which is associated with expected eyes-closed restful EEG. Also, the algorithm minimised the variance of the power spectrum and minimised instantaneous correlation (zero lag correlation) over all electrodes to eliminate major artifacts.

The stability of an individual’s FN was defined to be the consistency of their PLF networks across all epochs in a given acquisition within a specific frequency band, as measured using 2-D correlation. This is based on network stability as presented and used by Shrey et al.^18^ and Smith et al.^25^. As multiple FNs were required for the correlation analysis the entire 150 s acquisition was split into 30 s epochs each overlapping their neighbour by half the epoch size (15 s) (Fig. 2b). This means all of the data were used and 9 epochs were created. For each of the 9 epochs, a PLF network was calculated. The stability of each EEG acquisition was then quantified by the 2-D correlation between each pair of the 9 PLF networks. Therefore, each PLF network was correlated to the other 8 PLF networks resulting in 36 correlations. The median stability was then taken as representative of the stability of the FNs for that EEG acquisition. The stability of the FNs was calculated across the 40 hours CR for all four frequency bands.

### Statistical Analysis

The PVT performance of the low and high impairment groups was compared using a non-parametric permutation test (*n* = 10, 000 permutations, two-tailed), establishing a *p*-value for the difference between the mean of the two groups for each measure over three equally sized ranges roughly split into biological day CR1 ([DLMO − 11 h, DLMO]), biological night ([DLMO + 1 h, DLMO + 12 h]) and biological day CR2 ([DLMO + 13 h, DLMO + 24 h]).

After the graph metrics were calculated from the baseline acquisitions, they were correlated to the level of PVT impairment using Pearson’s correlation. In addition, the Pearson’s correlation between the baseline graph metrics and PVT performance across the CR was calculated.

Within the stability analysis, increases in stability from prior to the WMZ to during the WMZ were assessed using a paired Cohen’s *d* test. The mean stability for each participant during the 3 hours prior to the WMZ was compared to the 3 hours during the WMZ. Normality of the data was determined through the Kolmogorov-Smirnov test. The associated *p*-values for both the group level and within high and low impairment groups are the results of associated paired two-tail *t*-tests. Furthermore, non-parametric permutation tests (*n* = 10, 000 permutations, two-tailed) were used to compare the stability of high and low impairment groups by comparing the mean for each participant across 12 hour periods. Finally, the Spearman’s rank correlation between stability and std of RT from the PVT was calculated. At the individual level the timeseries for stability and the std of RT across the CR were normalised to have a mean of 0 and a standard deviation of 1, to ensure comparable scales.

Throughout the analysis the p-values were adjusted for multiple comparisons using the Benjamini-Hochberg method^26^ and *p* < 0.05 was considered a significant result.

## Results

### Psychomotor Vigilance Test

The timeseries for each of the four PVT measures when the participants were split by low and high impairment are given in Fig. 3, corresponding group level figures are given in Supplementary Fig. S3. The general trend for all participants was stable good performance until ∼ DLMO + 5 h when the PVT measures increase, reflecting an increase and greater variability in the RTs. The greater increase in the high impairment group is reflected by the results of the permutation tests (*p* ≤ 0.0001), which show a significant difference between the two impairment groups for all PVT measures when averaged over [DLMO + 1 h, DLMO + 12 h] and [DLMO + 13 h, DLMO + 24 h]. Also, a significant difference was found between the high and low impairment group for mean RT (*p* = 0.0130) and percentage of PVT lapses (*p* ≤ 0.0001) when averaged over [DLMO − 11 h, DLMO].

Note that, a decrease in all measures, especially in the high impairment group, was seen around DLMO + 21 h, with a local minimum occurring during the second WMZ, a time when the circadian rhythm maximally promotes wakefulness.

### PVT Impairment

To assess if a baseline EEG acquisition provides information about the impairment of the participant under sleep deprivation the mean node strength, clustering coefficient, characteristic path length and stability of each baseline network were correlated to the level of impairment in each of the four PVT measures. The Pearson’s correlation and associated *p*-values are given in Supplementary Table S2. There were five metrics that significantly correlated to PVT impairment (Fig. 4). Indeed for B_FA_ the std of RT had a significant negative correlation with mean node strength (*p* = 0.0042) and a significant positive correlation with characteristic path length (*p* = 0.0013). Furthermore, for B_-11_ the std of RT had a significant negative correlation with mean node strength (*p* = 0.0028), characteristic path length (*p* = 0.0013) and stability (*p* = 0.0263). This indicates that networks with a high mean node strength, low characteristic path length and high stability are more robust to the effects of sleep deprivation as measured by std of RT. The figures for the other combinations are presented in the Supplementary Fig. S4 - S11.

**Figure 4.**
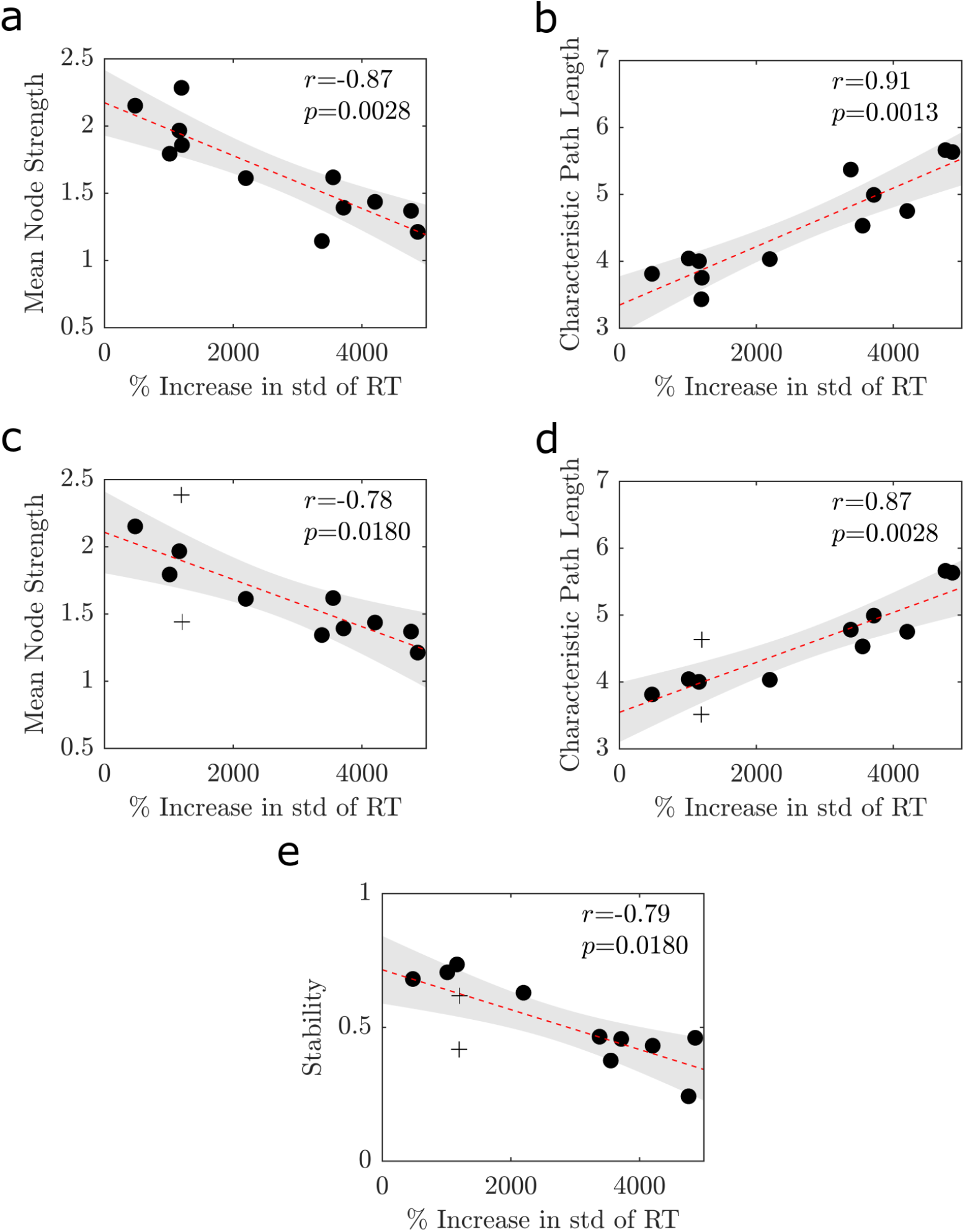
Significant Relationships Between PVT Impairment and Baseline Graph Metrics. Scatterplots relating the PVT impairment as measured using std of RT to **(a)** mean node strength for the first acquisition after awakening, **(b)** characteristic path length for the first acquisition after awakening, **(c)** mean node strength for the acquisition at DLMO − 11 h, **(d)** characteristic path length for the acquisition at DLMO − 11 h and **(e)** stability for the acquisition at DLMO − 11 h. For all plots, a black dot represents a participant, the red dashed line is the linear fit fitted using linear regression between the two variables and the grey patch represents the 95% confidence interval. In **(c**,**d**,**e)** the + represents the participants whose acquisitions at DLMO − 11 h differs from their first acquisition after awakening.

For the median RT two participants were considered outliers, as seen in Supplementary Fig. S4b. The analysis was repeated without these participants, however their removal did not greatly change the correlations as seen in Supplementary Table S3.

### PVT Performance

Within this section, the mean node strength, clustering coefficient, characteristic path length and stability of the baseline EEG acquisitions were correlated with PVT performance over the entire CR, rather than the single value used for PVT impairment. Note that positive correlations are indicative of high baseline graph metrics being associated with worse performance. The Pearson’s correlation between std of RT and the four graph metrics for B_FA_ are presented in Fig. 5. Complimentary figures for B_−11_ and the other PVT measures are in Supplementary Fig. S12 -S20.

**Figure 5.**
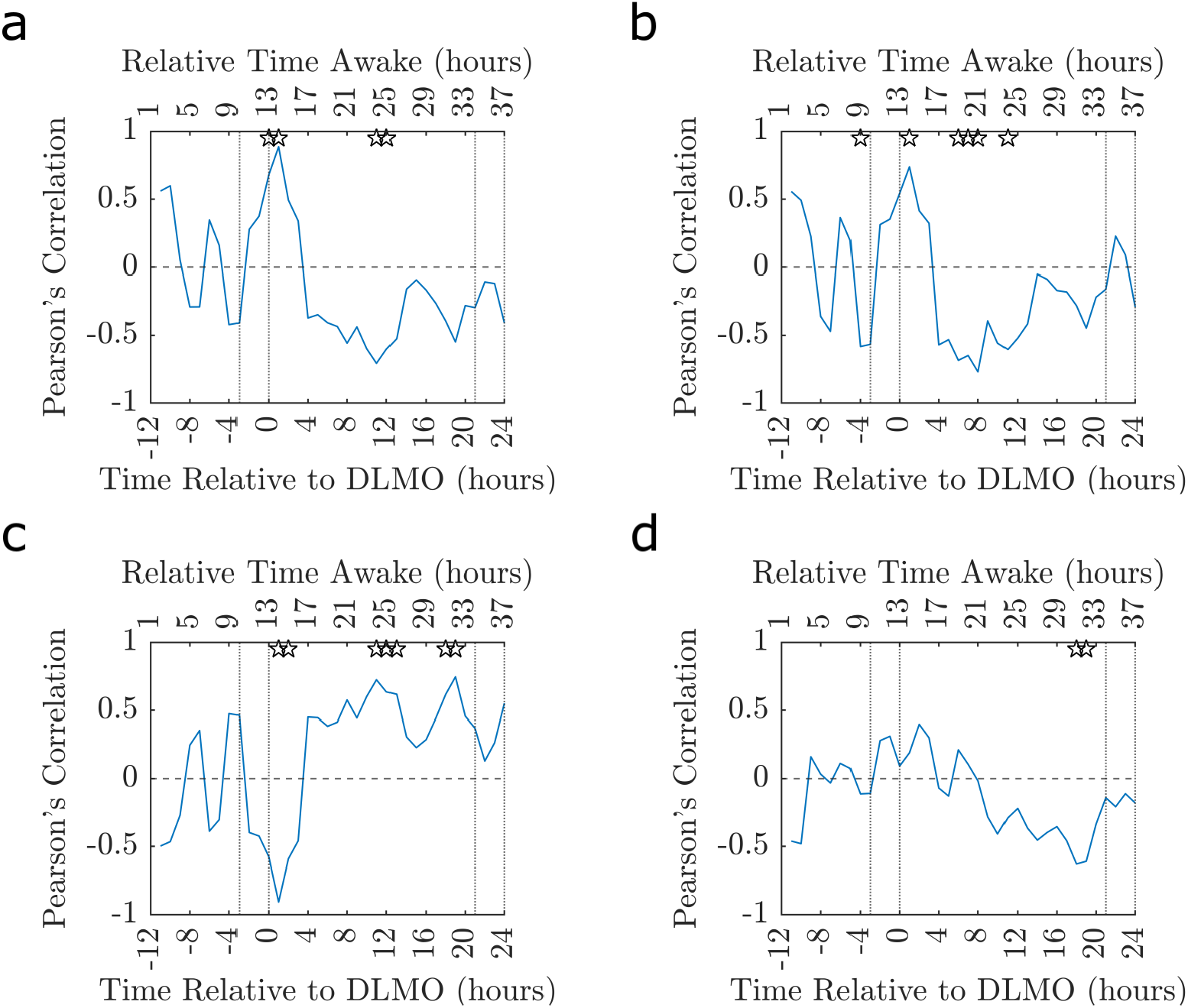
Pearson’s Correlation Between Baseline Graph Metrics for the First Acquisition and PVT Performance. How the Pearson’s correlation coefficient between baseline **(a)** mean node strength, **(b)** clustering coefficient, **(c)** characteristic path length and **(d)** stability for the first acquisition after awakening and the PVT performance as measured using std of RT. In all plots the ☆ indicates where the *p*-value associated with the correlation is significant *p* < 0.05 (uncorrected) and the dotted vertical lines indicate the times considered to be in the WMZ (3 hours before DLMO to 5 minutes after) and the relative time awake provides an indication of the time awake for the majority of the participants.

As seen in Fig. 5 the relationship between baseline graph metrics and PVT performance is more complex than the results presented from the PVT Impairment analysis suggest. There is a general trend that the correlation with maximum amplitude occurs at DLMO + 1 h before correlations with a slightly reduced amplitude and opposite sign occur near DLMO + 11 h. This change in correlation sign indicates that at DLMO + 1 h a high baseline mean node strength, clustering coefficient and stability as well as low characteristic path length are associated with poorer performance whilst near DLMO + 11 h these baseline network markers are associated with better performance. This inverse relationship is shown clearly through scatterplots in Supplementary Fig. S13. However, while the trend is very clear and fairly consistent across the 2 baseline acquisition, 4 graph metrics, 4 PVT performance metrics and 36 timepoints no significance remained after correcting the associated *p*-values for multiple comparisons.

### Stability

#### Group Level

The stability of each participant for a given EEG acquisition within a frequency band was calculated and Fig. 6 shows the group level stability across participants.

**Figure 6.**
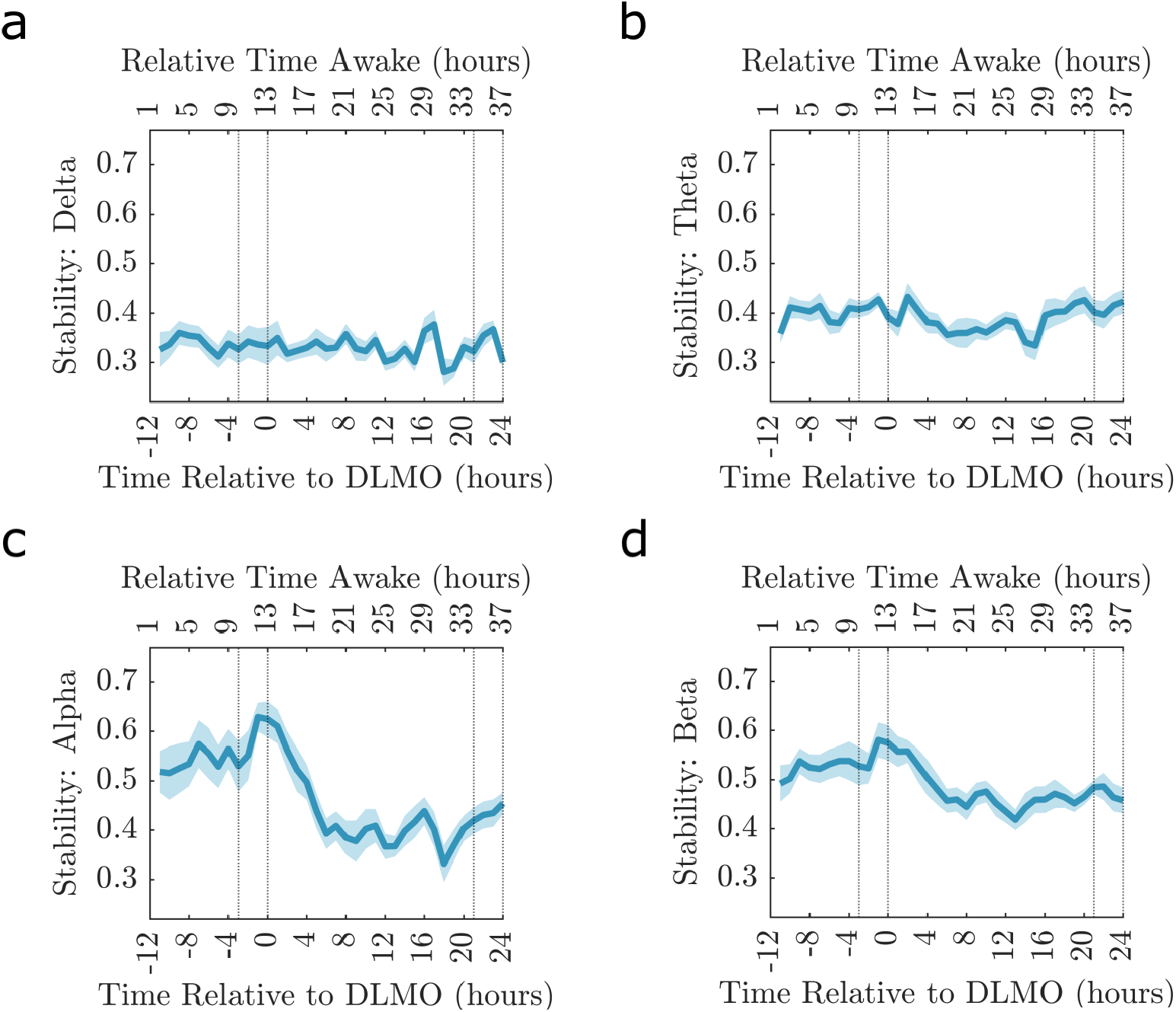
Stability of Participant’s FNs. The stability across all participants for a given frequency band **(a)** Delta, **(b)** Theta, **(c)** Alpha and **(d)** Beta. The solid blue line is the mean across the participants stability and the shaded area is the standard error of the mean. In all plots the dotted vertical lines indicate the times considered to be in the WMZ (3 hours before DLMO to 5 minutes after) and the relative time awake provides an indication of the time awake for the majority of the participants.

Both alpha and beta frequency bands show consistent levels of stability during the day before peaking in the WMZ. After the WMZ peak stability declined into the biological night before reaching a plateau, with some variation, during the second day. No visual trends were seen for Delta or Theta. Also, when comparing the stability prior to and during the first WMZ there were no significant differences for any frequency band as measured using permutation tests, as seen in Supplementary Table S4.

#### PVT Groups

High and low impairment groups based on std of RT were utilised for the stability analysis, due to the observation that FNs were associated with this PVT measure in both the PVT Impairment and Performance analysis.

As seen in Fig. 7 the two groups were significantly different during the first 12 h in the theta, alpha and beta bands (*p* = 0.0010, *p* < 0.0001 and *p* < 0.0001 respectively, non-parametric permutation test *n* = 10, 000), the middle 12 h in only the alpha band (*p* = 0.0030) and the final 12 h in the theta and alpha bands (*p* = 0.0010 and *p* < 0.0001, respectively). Indeed, in the alpha band, the low impairment group consistently had higher stability.

**Figure 7.**
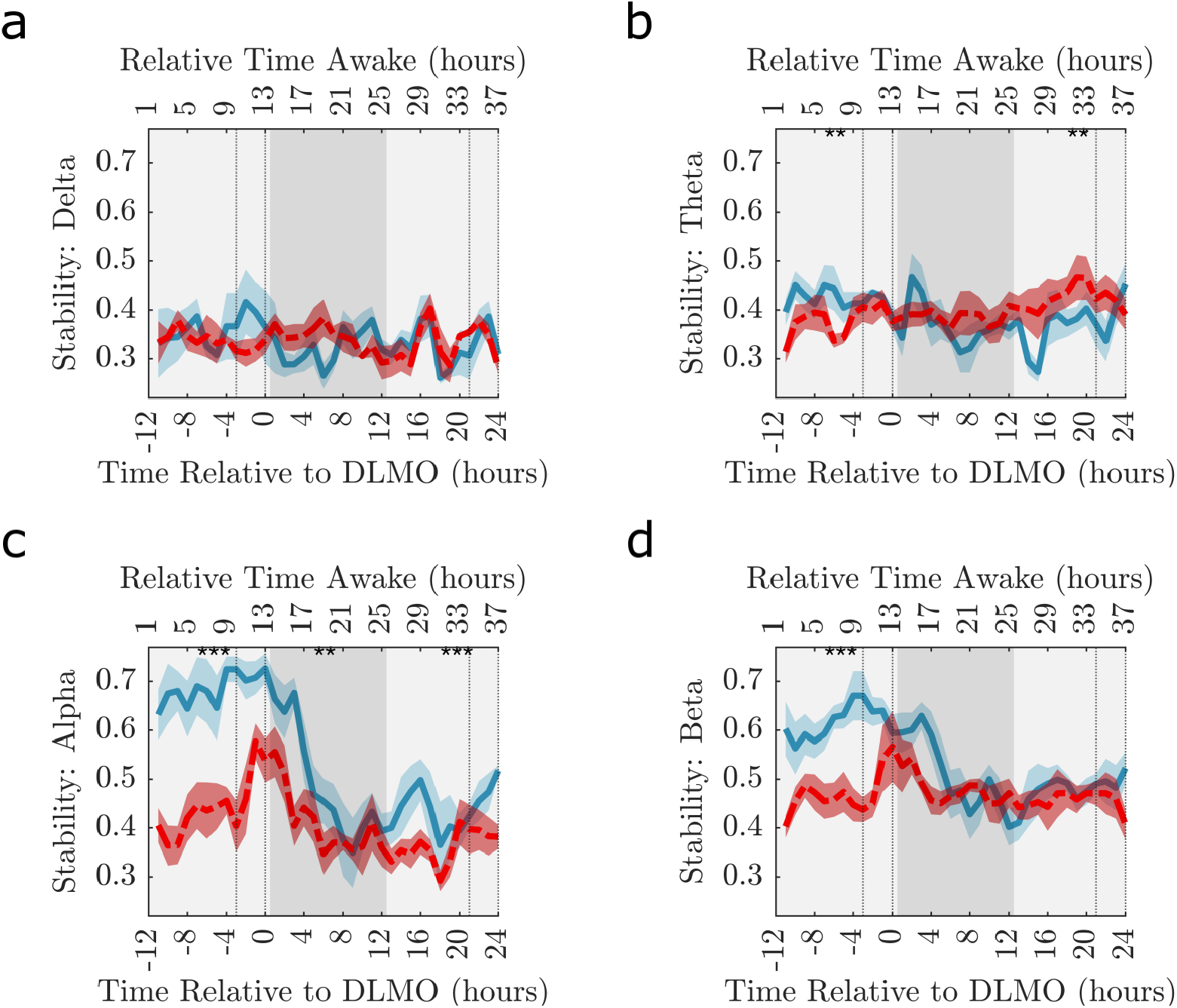
Stability of Participant’s FNs with Low and High Impairment. The stability across the participants with the lowest PVT impairment and the highest PVT impairment as measured using the percentage increase in standard deviation. The solid blue line 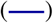 is the mean across the median stability of the 6 participants with the lowest impairment and the dashed red line 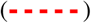 is the mean across the median stability of the 6 participants with the highest impairment. The corresponding blue and red shaded area is the standard error of the mean. Results are given for **(a)** Delta, **(b)** Theta, **(c)** Alpha and **(d)** Beta frequency bands. The significance of permutation tests (*n* = 10, 000) comparing the mean value of the low and high impairment groups over adjacent 12-hour period depicted by different shades of grey are represented by, * *p* < 0.05, ** *p* < 0.005, *** *p* < 0.0001 (*p*-values provided in Supplementary Table S5). In all plots the dotted vertical lines indicate the times considered to be in the WMZ (3 hours before DLMO to 5 minutes after) and the relative time awake provides an indication of the time awake for the majority of the participants.

When comparing the stability prior to and during the first WMZ no significant differences occurred for any frequency band after correcting for multiple comparisons as seen in Table 1.

**Table 1.**
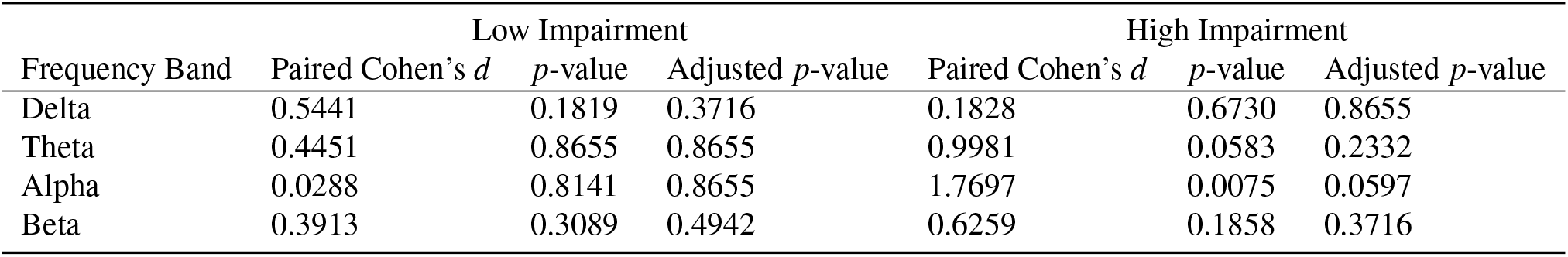
Stability Prior to and During the WMZ for Groups of High and Low Impairment. Paired Cohen’s *d* and the associated *p*-values for paired *t*-tests comparing participants mean stability 3 hours prior to the WMZ to the mean stability during the 3 hours of the WMZ for a given frequency band within the high and low impairment group. Both uncorrected values and *p*-values corrected for multiple comparisons using the Benjamini-Hochberg method^26^.

#### Individual Level

To directly understand the relation between std of RT over the CR and the stability of the participant’s network these time-series were also compared using Spearman’s Rank correlation, this is displayed in Supplementary Fig. S21 and Supplementary Table S6.

Standard deviation of RT generally worsened over time and stability decreased, as summarised through the Spearman’s Rank Correlations of −0.4448 ± 0.2876 (mean ± std) and 8/13 participants having a significant inverse relationship. Indeed, only Participant 12 had a positive, albeit non-significant, correlation, likely due to their stability showing no noticeable drop below the first acquisition, unlike the other participants.

## Discussion

Within this study, we sought to understand the relationship between sleep deprivation and functional networks in the brain. In particular, we used only information from baseline EEG acquisitions to explore associations with robustness to sleep deprivation. Graph metrics from a baseline EEG acquisition were associated with the level of impairment, defined as the magnitude of sleep deprivation effects irrespective of time, on the PVT. High mean node strength, low characteristic path length and high stability at baseline significantly correlated with lower levels of impairment as measured using PVT (Fig. 4). This shows that a significant association between baseline FNs and task performance under sleep deprivation exists. A natural extension was to consider the association between baseline EEG acquisitions and PVT performance across the CR. This analysis showed that high mean node strength, high clustering coefficient, low characteristic path length and high stability at baseline were associated with more resilience to sleep deprivation during the second day of the CR, as measured using PVT RTs. Furthermore, these graph metric features were more strongly associated with reduced performance at DLMO + 1 h - when the drive to sleep starts to increase. Hence, in general the performance of the group decreased with time awake but within the group the ranking of the participants changed such that the individuals with the worst performance at DLMO + 1 h where among the best performers at DLMO + 11 h, as seen in Supplementary Fig. S13.

Pearson’s correlation using characteristic path length for both PVT impairment and performance demonstrated a similar trend to mean node strength, clustering coefficient and stability, however with the signs reversed. The association between high baseline mean node strength and better PVT performance after longer exposure to sleep deprivation is analogous to the results of Mu et al.^27^ and Caldwell et al.^28^ who both found associations between higher levels of global brain activity, as measured using fMRI, with resilience to the effects of sleep deprivation. Furthermore, when combining the results for the clustering coefficient and characteristic path length this suggests that participants with a small-world network at baseline are more vulnerable to sleep deprivation at the point of biological sleep onset immediately after DLMO. However, a baseline small-world network results in participants being more robust to sleep deprivation after being exposed for a longer period of time. Overall, this suggests that a more efficient and stable baseline network results in increased vulnerability to sleep near sleep onset and increased resilience to sleep near the end of the biological night. Potentially this could reflect the ability of the network to efficiently propagate the signals for sleep and wakefulness triggered by the circadian rhythm.

The substantial negative correlations between baseline mean node strength and clustering coefficient and std of RT near [DLMO + 8 h, DLMO + 12 h] and positive correlations between baseline characteristic path length and std of RT near DLMO + 12 h (Fig. 5) are consistent with the results from the PVT Impairment analysis, as seen in Fig. 4, although the magnitude is lower. Furthermore, the oppositely signed peak at DLMO + 1 h was typically higher in amplitude than the correlations from the impairment analysis. Within this study we aligned the data by DLMO and know that the drive to wakefulness in the WMZ and subsequent drive to sleep post DLMO is closely aligned across participants. Indeed, we know that performance variability across participants is low during the WMZ and increases further away from DLMO^9^. Therefore, at DLMO + 1 h less inter-individual differences in the timing of circadian and homeostatic process could have contributed to the greater effect size at DLMO + 1 h. Correspondingly, the increased variability in PVT performance across participants near [DLMO + 8 h, DLMO + 12 h] and intrinsic variability in the timing of each participants worst performance (Supplementary Table S1) could have contributed to the reduced effect size for the performance analysis compared to the impairment analysis.

Baseline EEG derived graph metrics do not correlate with PVT performance measures uniformly across the CR, with a switch occurring around DLMO into the biological night. As the baseline network for a participant remains constant this change in correlation is driven by changes in the participants’ PVT performance. Indeed, while sleep deprivation has a general negative effect on performance, the ranking of the participants almost reverses (Supplementary Fig. S13) such that the best individual for a task at DLMO + 1 h would actually be among the worst to complete the task at DLMO + 11 h. Slight changes in the ranking over time would be expected due to inter-individual variability, however the reverse in the PVT performance ranking could be due to the influence of a participants’ circadian strength (the amplitude of an individual’s circadian rhythm). This is because, if the monotonically increasing homeostatic process were the only influence, then the order of sleep deprivation susceptibility within the group would have no reason to reverse. However, a strong circadian strength that pushes an individual more towards sleep at night (worst performance) but also more towards alertness in the day (best performance) could explain reversed rankings and hence the change in correlation sign. We theorise that this change in the Pearson’s correlation sign could be a result of the strength of circadian rhythm and hence the baseline graph metrics may be associated with the circadian strength. However, circadian strength is a relatively new area of research and has not been extensively studied. Currently, markers of circadian strength are measured using core body temperature or brain temperature. There is evidence that brain temperature is related to neurological conditions including epilepsy and mood disorders^29^ and circadian changes in brain temperature have been found to be predictive of survival after brain injury, with the absence of circadian changes indicative of lower survival rate^30^. Consequently, future studies incorporating brain temperature measurements, preferably using non-invasive approaches such as magnetic resonance spectroscopy^31^, are needed to fully understand the connection between graph metrics and circadian strength.

The std of RT encompasses both the sporadic good RTs and lapses, that are consistent with the state instability hypothesis^8^, potentially explaining why these correlations were consistently high across the impairment and performance analysis. Indeed, Van Dongen et al.^4^ suggests that state instability could be due to parts or even the whole brain falling asleep for brief periods of time, neighboured by periods of normal wakefulness - note only regional sleep could occur within this CR protocol. Hence, future work could assess the association between regional sleep in the brain and std of RT. Furthermore, as the median, by the nature of the metric, removes extreme RTs, this could explain why the measure consistently has the weakest correlations.

Throughout this analysis the graph metrics were calculated using FNs created using the alpha frequency band. The alpha band was chosen due to the literature which shows the alpha band has the highest spectral content during the baseline acquisitions^32^. Also, Chua et al.^6^ found that only the alpha power spectrum could explain significant amounts of the variance between participants when they were sleep deprived. However, extending this research to include additional frequency bands could be of interest. Furthermore, while global graph metrics were considered in this study the level of correlation for individual electrodes could be considered in the future. For instance, Verweij et al.^33^ found that only scalp electrodes placed above the prefrontal and frontal brain regions were significantly impacted by sleep deprivation, therefore a focus on these electrodes could result in greater correlations.

Furthermore, motivated by the instability hypothesis^8^ the stability of the FNs across the CR was calculated, to understand how the brain reacts to sleep deprivation both in the long term over the protocol and the short term within an EEG acquisition. The stability across all participants in the alpha and beta frequency bands (Fig. 6) demonstrated an inverse relation to the PVT measures. Importantly, the stability of the FNs showed peaks during the WMZ, highlighting the effect of the circadian rhythm, while the drop in stability during the biological night shows the influence of the homeostatic pressure. This corresponds to the steep decrease in PVT performance during the biological night ∼ 5 hours after DLMO before a marked improvement during the WMZ, as seen in Fig. S3. The similar relation between stability and PVT performance could provide evidence for the state instability hypothesis^8^, which suggests that brief returns to baseline levels occur alongside a general decrease in performance creating more variability. Indeed, instability and std of RT both increase with sleep deprivation reflecting increased variability in both FNs and behavioural performance, respectively.

When the participants were split into impairment groups, we saw a stark difference in their stability for the theta, alpha and beta bands during the first 12 hours, despite this being a time when neither group would have been experiencing sleep deprivation. When combined with the lack of significance difference between the high and low impairment groups for the median and std of RT (Fig. 7) this could suggest that the level of impairment a participant faces when sleep deprived is due to a mechanism that is present throughout the day, but only significantly affects performance when sleep debt accumulates. Interestingly, it is through sleep deprivation that the stability of the groups becomes more similar. This occurs due to the low impairment group decreasing their level of stability, rather than any marked change in the high impairment group. It is worth noting that the alpha band maintains a significant difference across the CR (*p* ≤ 0.0032). When combined with the PVT performances significantly differing (*p* < 0.0001) across all four measures during the second and third 12 hours of the protocol, Fig. 3, this could suggest that the stability in the alpha band is the most informative and most closely linked to PVT performance, as also indicated by the work of Chua et al.^6^.

Our results for stability could provide evidence to support a link between excitability and stability. For example, in the alpha and beta bands, especially for the low impairment group, stability generally decreases with increased time awake whilst excitability is known to increase with sleep deprivation^34,35^. Therefore, this suggests that high stability could be interpreted as less excitability and vice versa. Furthermore, we saw a visual increase in stability in the alpha and beta frequency bands as participants enter the first WMZ (Fig. 6) - especially in the high impairment group (Fig. 7) and this was also visually seen during the second WMZ for the low impairment group. Ly et al.^34^ found a reduction in cortical excitability from prior to during the WMZ, as measured by the significant decrease in the amplitude of a TMS-evoked EEG potential in the supplementary motor area. Again supporting the link between lower excitability and higher stability. Why the increase in stability during the WMZ is seen for different impairment groups in the two WMZs could be the focus of future studies.

At the individual level 8 participants showed a significant association between stability of the baseline FNs and PVT performance (Supplementary Fig. S21). For instance, Participant 8 exhibited a clear inverse relationship between the stability of the network and the std of RT. The variability in the Spearman’s correlation between PVT performance and stability appears to be linked to the impairment groups as those that are the least impaired see a decrease in their stability over time, which more strongly corresponds to an increase in their std of RT. While for some participants a link between stability of FNs and the stability of their PVT RTs is seen the cause of this relationship remains unclear. However, it in unlikely that the stability of the FN directly impacts task performance, rather the same mechanism that impacts task performance also manifests as a less robust FN. For instance, prolonged wakefulness is known to increase the concentration of excitatory neurotransmitters^36^. Future research could investigate if there is an underlying link between neurotransmitters and the stability of FNs or alternatively PVT performance.

A natural extension of this research would be the prediction of individual susceptibility to sleep deprivation, using baseline graph metrics. Understanding susceptibility without requiring sleep deprivation could offer vital insights into the work patterns suitable for employees, especially in careers where acute sleep deprivation and sleep debt are likely and even unavoidable. For example, in the US 40.1% of healthcare professionals report getting insufficient sleep defined as less than 7 h of sleep while only 33% of military service members reported insufficient sleep, defined as < 7 h for adults >22 years and < 8 h for adults <21 years^37,38^. With further development, individualised prediction of sleep deprivation vulnerability could be used in a targeted manner within the employment process to help prioritise people for additional resources, mitigating their increased impairment before incidents occur.

In addition, throughout the correlation analysis at most two EEG acquisitions were considered per person. It is reasonable that other EEG acquisitions over the 40 h CR would have high levels of association and could be the focus of future research, especially as Chua et al.^6^ created one predictor using data across the first 16 h of wakefulness. However, our analysis could be seen as streamlining the approach of Chua et al.^6^ as high effect sizes (*r* = 0.89) from a singular EEG acquisition were found, increasing practically for any real world application.

The results should be interpreted with the following limitations in mind. First, the PVT may not reflect actual vulnerability to sleep deprivation with respect to real-world operations, despite its utility in sleep deprivation research^14^. While it has been associated with memory^39^, lane position and speed variability in a simulated driving scenario^40^ as well as mathematical fluency and reading comprehension in children^41^ future work should examine how well PVT markers reflect performance in other tasks.

Second, for the PVT analysis the dataset included 13 participants and thus we urge caution with the generalisabililty of the results. However, these 13 participants underwent a long and demanding protocol which resulted in large amounts of data from each individual, leading to significant correlations and differences. Hence, motivating further studies with additional participants.

Third, the participants in this study were selected to create a homogeneous group. This is typical in sleep deprivation and biomarker development studies to reduce individual differences^42^. Indeed, this study only included two females. This may be problematic due to the clear sex difference in response to sleep loss^43–45^. However, to minimise any sex difference effect, women were tested in the follicular phase of their cycle, when they look, from a performance perspective, closer to men^45^. Beyond sex, age also modulates the response to sleep loss such that older adults were found to be less impacted by sleep deprivation, as measures using PVT, compared to younger adults^44^. As this study utilised young adults, our results may not generalise to older populations. In addition, Facer-Childs et al.^46^ found that PVT performance profiles differ between extreme chronotypes and are significantly different at 08: 00, which is similar to the time of both baseline acquisitions. Taken together, the framework presented in this paper could be used to investigate differences in the associate of FNs and sleep deprivation across sex, age and chronotype differences.

Overall, we have considered the problem of sleep deprivation from two angles: how are baseline FNs associated with the effect of sleep deprivation and how does the stability of the FNs help to understand the effect of sleep deprivation? We have seen that lower mean node strength, lower clustering coefficient, higher characteristic path length and less stability are features strongly associated with greater levels of impairment after a night of sleep deprivation, as measured using std of RT. However, these same features are associated with improved performance at DLMO + 1 h. Therefore, whilst the performance generally decreases throughout the CR the ranking of the participant’s performance almost reversed, suggesting the optimum person to complete a task at DLMO + 1 h is actually amongst the worst at DLMO + 11 h, highlighting the time dependent nature of this analysis. The topology of the FNs could be related to a combination of circadian strength and the ability to propagate signals related to the circadian rhythm. Also, the stability of the network may help to explain why some participants are more resilient to the effects of sleep deprivation and why PVT performance generally decreases with wakefulness.

## Acknowledgements

Sophie L. Mason acknowledges the financial support of the University of Birmingham through an Alumni Scholarship and the N-CODE Network+ funded by Engineering and Physical Sciences Research Council (EPSRC)/ Medical Research Council (MRC) via grant EP/W035030/1. Leandro Junges acknowledges support from a Waterloo Foundation research grant (Ref no. 1970-4687) and from the University of Birmingham Dynamic Investment Fund. Wessel Woldman acknowledges the financial support of Epilepsy Research UK through an Emerging Leader Fellowship (F2002) and from the University of Birmingham Dynamic Investment Fund. Andrew P. Bagshaw acknowledges EPSRC (EP/J002909/1), an Institutional Strategic Support Fund Accelerator Fellowship from the Wellcome Trust (Wellcome, 204846/Z/16/Z) and BBSRC funding (BB/X013634/1). John R. Terry acknowledges the financial support of UKRI (EPSRC) via Fellowship EP/T027703/1 and Network+ grant EP/W035030/1 and from the University of Birmingham Dynamic Investment Fund. The publication of this article is supported by the University of Birmingham’s Open Access Fund and by a Project Grant from the Cooperative Research Centre for Alertness, Safety, and Productivity, Melbourne, Australia (1.1.01_15).

The authors also wish to acknowledge the technicians, staff and students of the Monash University Sleep and Circadian Medicine Laboratory who aided data collection. We also thank Drs. Leilah Grant, William McMahon and Jinny Collet for assistance in collecting data in the study; Caroline Beatty for recruitment support; study physicians Drs. Simon Joosten and Eric Kuo; Dr Sylvia Nguyen for conducting psychiatric screening; Dr Charmaine Diep for assisting in data transfer; Kellie Hamill for logistical help and Professor Shantha Rajaratnam and Professor Steven Lockley for assistance in the design of the larger study. We also gratefully acknowledge the assistance of the Adelaide Research Assay Facility for providing plasma melatonin analysis.

## Author contributions

Conception and Design: SLM, LJ, JRT, APB, CA, SF. Data Collection: CA, SF. Data Analysis and Interpretation: SLM, LJ, WW, JRT, APB, CA. Drafting and Revising the Manuscript: All authors.

## Data Availability Statement

The datasets analyzed during the current study are available upon reasonable request.

## Additional information

WW and JRT are co-founders of Neuronostics Ltd. CA reports no conflicts of interests related to the data or results reported in this paper. In the interest of full disclosure: CA has received a research award/prize from Sanofi-Aventis; contract research support from VicRoads/Department of Transport Victoria, Rio Tinto Coal Australia, National Transport Commission, Tontine/Pacific Brands, Transport Accident Commission, Takeda Pharmaceutical; and lecturing fees from Brown Medical School/Rhode Island Hospital, Ausmed, Healthmed and TEVA Pharmaceuticals; and reimbursements for conference travel expenses from Philips Healthcare. In addition, she has served as a consultant through her institution to the Rail, Bus and Tram Union, the Transport Accident Commission and the National Transportation Committee. She has also served as an expert witness and/or consultant in relation to fatigue and drowsy driving. CA was a Theme Leader in the Cooperative Research Centre for Alertness, Safety and Productivity. SF reports no conflicts of interests related to the data or results reported in this paper. In the interest of full disclosure: SF reports receiving consultancy fees from Circadian Therapeutics and AMO Pharma. All other authors declared that the research was conducted in the absence of any commercial or financial relationships that could be construed as a potential conflict of interest.

