## Supplementary Material for "Associating EEG Functional Networks and the Effect of Sleep Deprivation as Measured Using Psychomotor Vigilance Tests"

#### Data Available

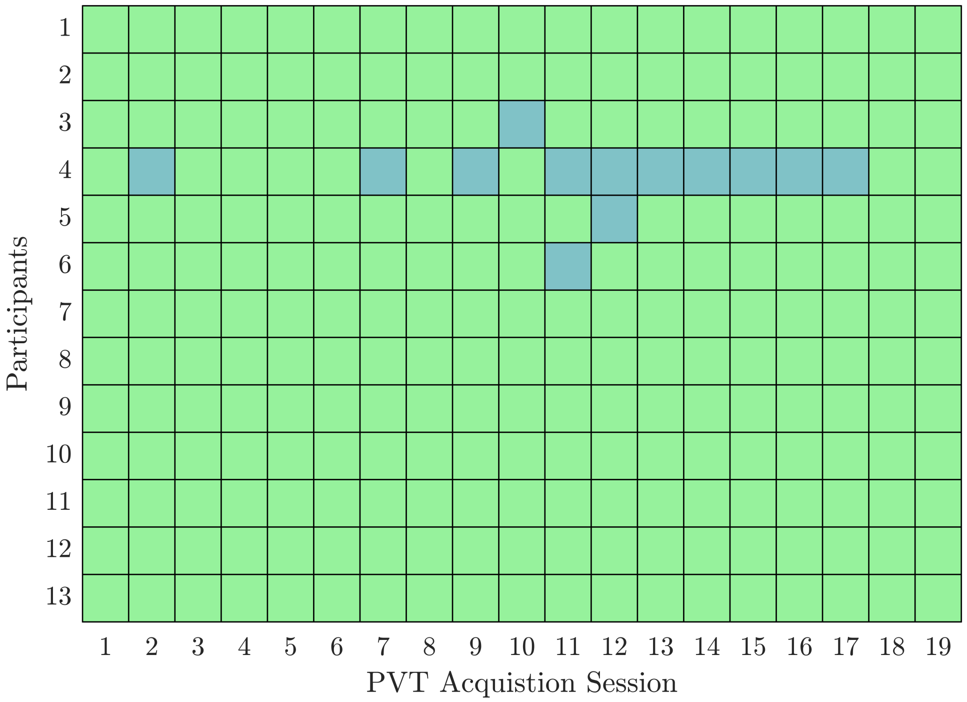


**Figure S1. PVT Acquisitions Available.** The available PVT acquisition sessions for each participant across the CR. The colours represent [green] a successful PVT acquisition and [blue] unable to record a PVT acquisition.

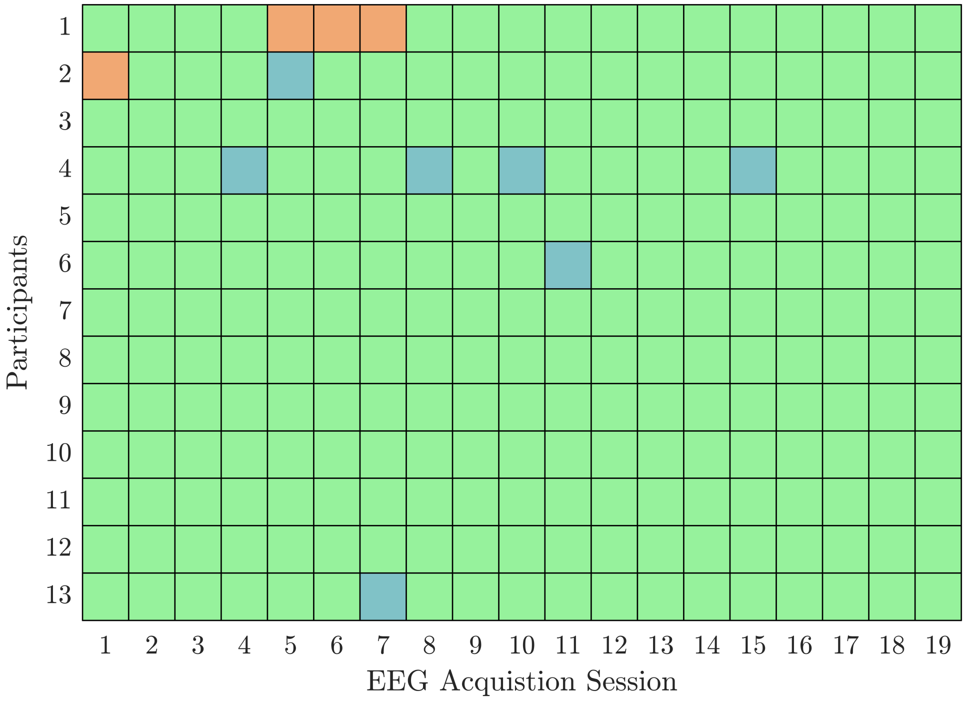


**Figure S2. EEG Acquisitions Available.** The available EEG acquisition sessions for each participant across the CR. The colours represent a successful EEG acquisition, an EEG acquisition with at least one faulty electrode and unable to record an EEG acquisition.

#### PVT Impairment and Performance

| Participant | Mean RT (ms) | | Median RT (ms) | | Standard Deviation of RT (ms) | | Percentage of Lapses | |
|---|---|---|---|---|---|---|---|---|
| 1 | -1,0 | 17,18 | -11,-10 | 9,10 | -3,-2 | 15,16 | - | 9,10 |
| 2 | -2,-1 | 14,15 | 0 | 20,21 | 0 | 14,15 | - | 20,21 |
| 3 | -2,-1 | 10,11 | -2,-1 | 10,11 | -2,-1 | 12,13 | - | 10,11 |
| 4 | -4,-3 | 6,7 | -4,-3 | 6,7 | -4,-3 | 6,7 | - | 6,7 |
| 5 | -11 | 20,21 | -6,-5 | 20,21 | 0 | 20,21 | - | 20,21 |
| 6 | -3,-2 | 9,10 | -3,-2 | 6,7 | -1,0 | 9,10 | - | 6,7 |
| 7 | -9,-8 | 19,20 | -9,-8 | 23,24 | -9,-8 | 19,20 | - | 23,24 |
| 8 | -10,-9 | 10,11 | -11 | 10,11 | -10,-9 | 10,11 | - | 10,11 |
| 9 | -1,0 | 9,10 | -1,0 | 17 | -1,0 | 9,10 | - | 15,16 |
| 10 | -2,-1 | 8,9 | -2,-1 | 8,9 | 0 | 16,17 | - | 8,9 |
| 11 | -7,-6 | 11,12 | -7,-6 | 21,22 | -7,-6 | 11,12 | - | 11,12 |
| 12 | -9,-8 | 17,18 | -9,-8 | 19,20 | -9,-8 | 17,18 | - | 19,20 |
| 13 | -2 | 10,11 | -6,-5 | 10,11 | -1,0 | 10,11 | - | 10,11 |

**Table S1. Timings of the Minimum and Maximum PVT Performance with Respect to DLMO.** For each of the four PVT performance measures the timing of the minimum mean RT, median RT, standard deviation of RT and percentage of lapses with respect to DLMO is given in the first column and the maximum is given in the second column. The timing for the minimum percentage of lapses is not provided as this is not used when assessing impairment.

#### PVT Group Level

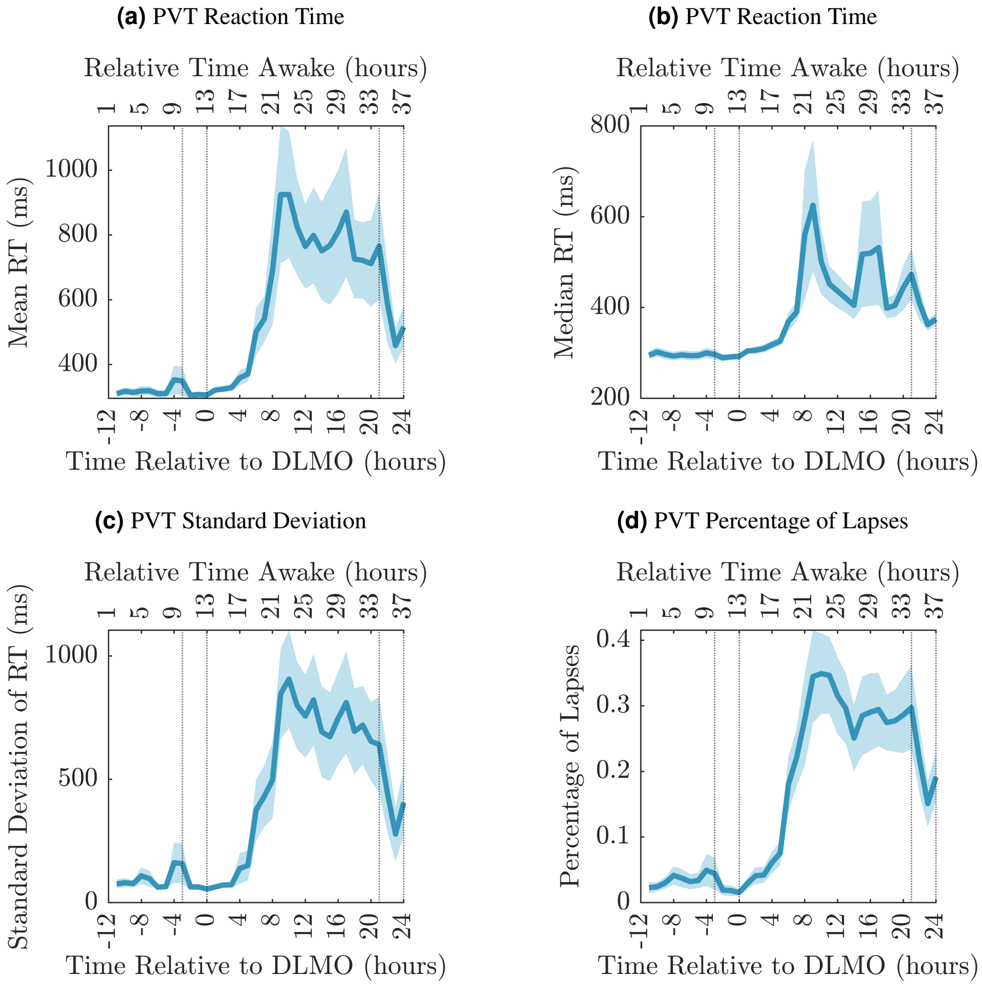


**Figure S3.** **Psychomotor Vigilance Test Performance.** The performance of all participants in the PVT as measured using **(a)** mean RT, **(b)** median RT, **(c)** standard deviation of RT and **(d)** percentage of lapses at each PVT session, relative to DLMO (time zero). The solid blue line is the mean across the participants' PVT metric and the shaded area is the standard error of the mean. The dotted vertical lines indicate the times considered to be in the WMZ (3 hours before DLMO to 5 minutes after) and the relative time awake provides an indication of the time awake for the majority of the participants.

#### PVT Impairment

| Baseline Acquisition: Graph Metric: PVT Impairment | Correlation | $p$-value | Adjusted $p$-value |
|---|---|---|---|
| $B_{FA}$: $\langle s \rangle$: Mean RT | -0.5618 | 0.0573 | 0.1529 |
| $B_{FA}$: $\langle s \rangle$: Median RT | -0.3538 | 0.2592 | 0.3054 |
| $B_{FA}$: $\langle s \rangle$: std RT | -0.8667 | 0.0003 | 0.0028* |
| $B_{FA}$: $\langle s \rangle$: % Lapses | -0.3748 | 0.2299 | 0.3054 |
| $B_{FA}$: $\langle C \rangle$: Mean RT | -0.4315 | 0.1613 | 0.2986 |
| $B_{FA}$: $\langle C \rangle$: Median RT | -0.3669 | 0.2407 | 0.3054 |
| $B_{FA}$: $\langle C \rangle$: std RT | -0.6078 | 0.0360 | 0.1441 |
| $B_{FA}$: $\langle C \rangle$: % Lapses | -0.3516 | 0.2624 | 0.3054 |
| $B_{FA}$ :$\lambda_L$: Mean RT | 0.6199 | 0.0315 | 0.1441 |
| $B_{FA}$ :$\lambda_L$: Median RT | 0.4171 | 0.1773 | 0.2986 |
| $B_{FA}$ :$\lambda_L$: std RT | 0.9093 | <0.0001 | 0.0013* |
| $B_{FA}$ :$\lambda_L$: % Lapses | 0.4605 | 0.1320 | 0.2901 |
| $B_{FA}$: $r_s$: Mean RT | -0.4022 | 0.1950 | 0.2986 |
| $B_{FA}$: $r_s$: Median RT | -0.2351 | 0.4619 | 0.4619 |
| $B_{FA}$: $r_s$: std RT | -0.5662 | 0.0550 | 0.1529 |
| $B_{FA}$: $r_s$: % Lapses | -0.2412 | 0.4502 | 0.4619 |
| $B_{-11}$: $\langle s \rangle$: Mean RT | -0.5490 | 0.0645 | 0.1588 |
| $B_{-11}$: $\langle s \rangle$: Median RT | -0.3520 | 0.2618 | 0.3054 |
| $B_{-11}$: $\langle s \rangle$: std RT | -0.7793 | 0.0028 | 0.0180* |
| $B_{-11}$: $\langle s \rangle$: % Lapses | -0.4327 | 0.1600 | 0.2986 |
| $B_{-11}$: $\langle C \rangle$: Mean RT | -0.3419 | 0.2768 | 0.3054 |
| $B_{-11}$: $\langle C \rangle$: Median RT | -0.2817 | 0.3750 | 0.4000 |
| $B_{-11}$: $\langle C \rangle$: std RT | -0.4085 | 0.1874 | 0.2986 |
| $B_{-11}$: $\langle C \rangle$: % Lapses | -0.3524 | 0.2612 | 0.3054 |
| $B_{-11}$ :$\lambda_L$: Mean RT | 0.6566 | 0.0204 | 0.1086 |
| $B_{-11}$ :$\lambda_L$: Median RT | 0.4563 | 0.1360 | 0.2901 |
| $B_{-11}$ :$\lambda_L$: std RT | 0.8750 | 0.0002 | 0.0028* |
| $B_{-11}$ :$\lambda_L$: % Lapses | 0.5741 | 0.0510 | 0.1529 |
| $B_{-11}$: $r_s$: Mean RT | -0.5843 | 0.0460 | 0.1529 |
| $B_{-11}$: $r_s$: Median RT | -0.3465 | 0.2699 | 0.3054 |
| $B_{-11}$: $r_s$: std RT | -0.7854 | 0.0025 | 0.0180* |
| $B_{-11}$: $r_s$: % Lapses | -0.4014 | 0.1960 | 0.2986 |

**Table S2. Results of correlating the Baseline Graph Metric with PVT Impairment.** The Pearson's correlation between the participants' PVT impairment and the graph metric for both baseline acquisitions when correlating mean node strength ($\langle s \rangle$), clustering coefficient ($\langle C \rangle$), characteristic path length ($\lambda_L$) and stability ($r_s$) with the four PVT measures. The $p$-values were corrected for multiple comparisons using the Benjamini-Hochberg correction for FDR[1]. * denotes significance $p$-value < 0.05.

#### Mean Node Strength

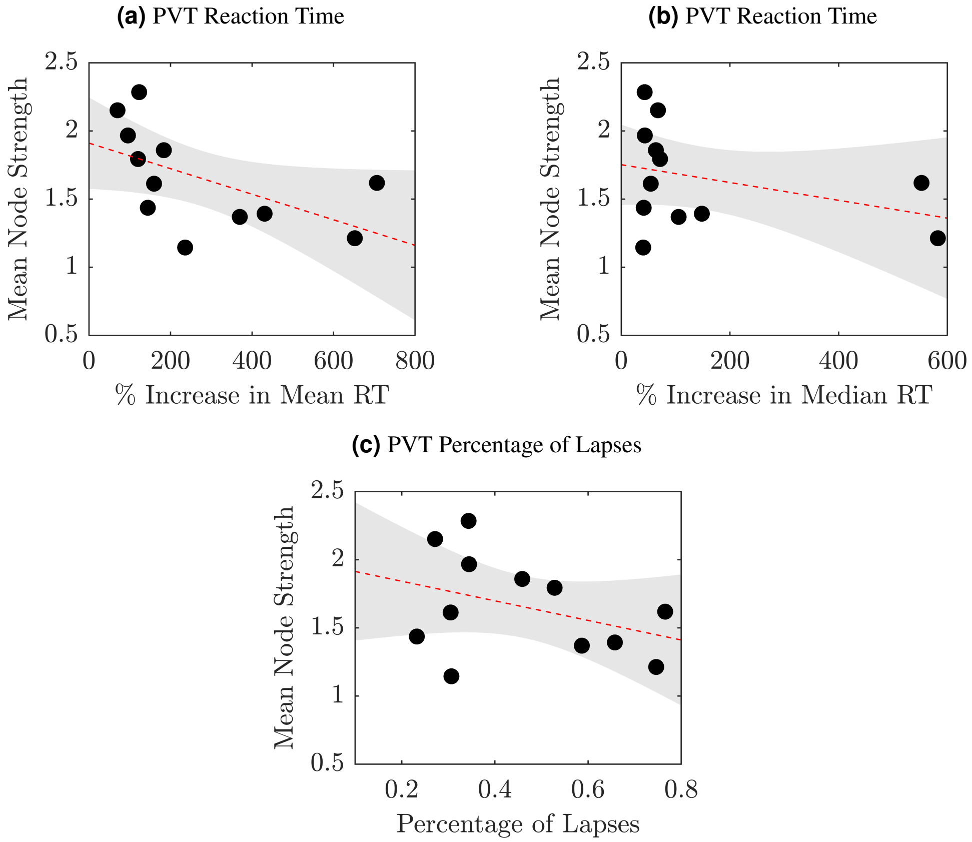


**Figure S4. Scatterplot Relating PVT Impairment to Mean Node Strength for the First Acquisition After Awakening.** Scatterplots relating the PVT impairment as measured using **(a)** mean RT, **(b)** median RT and **(c)** percentage of lapses to mean node strength for the first acquisition after awakening. For all plots, a black dot represents a participant, the red dashed line is the linear fit fitted using linear regression between the two variables and the grey patch represents the 95% confidence interval.

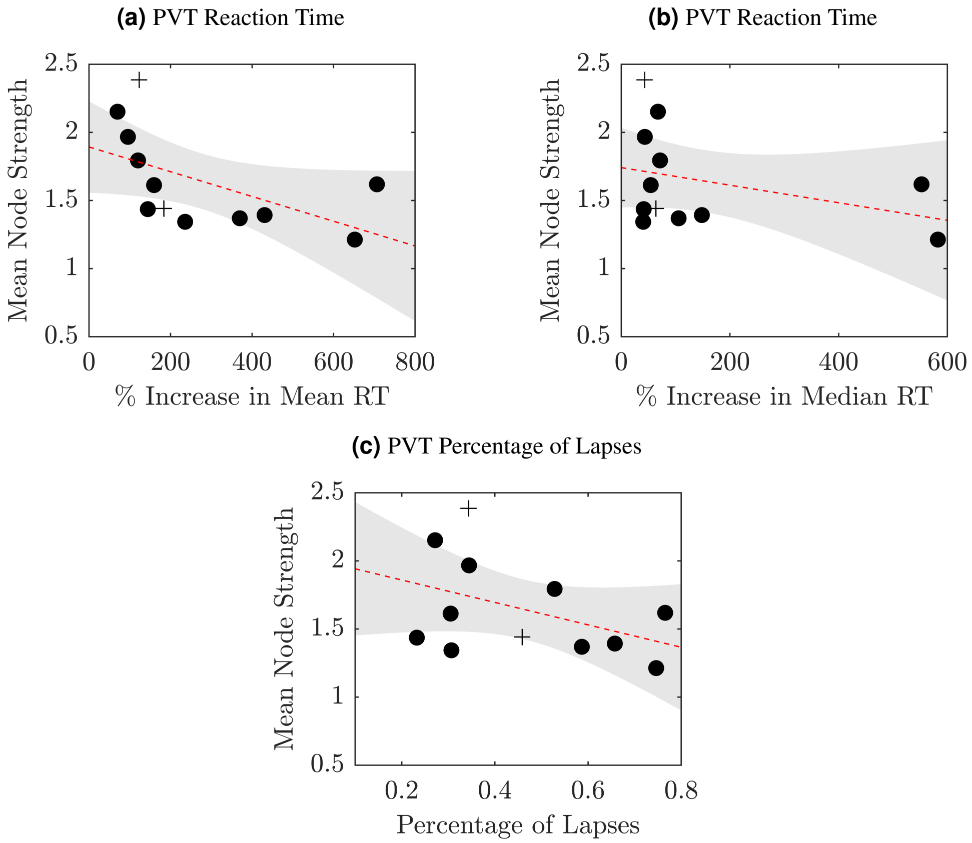


**Figure S5. Scatterplot Relating PVT Impairment to Mean Node Strength for the Acquisition 11 Hours Before DLMO.** Scatterplots relating the PVT impairment as measured using **(a)** mean RT, **(b)** median RT and **(c)** percentage of lapses to mean node strength for the acquisition 11 hours before DLMO. For all plots, participants whose EEG acquisition 11 hours before DLMO is not their first recording are represented by +, while those whose first EEG acquisition is 11 hours before DLMO are denoted by • a black dot, the red dashed line is the linear fit fitted using linear regression between the two variables and the grey patch represents the 95% confidence interval.

#### Clustering Coefficient

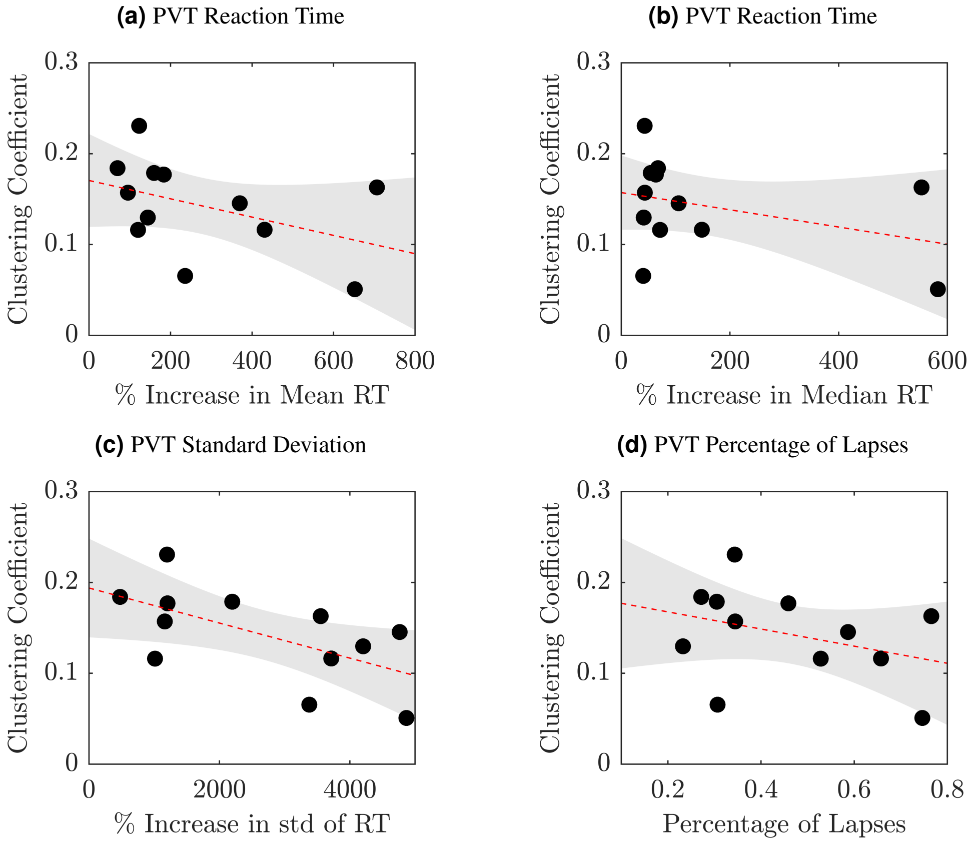


**Figure S6. Scatterplot Relating PVT Impairment to the Clustering Coefficient for the First Acquisition After Awakening.** Scatterplots relating the PVT impairment as measured using **(a)** mean RT, **(b)** median RT, **(c)** standard deviation of RT and **(d)** percentage of lapses to the clustering coefficient for the first acquisition after awakening. For all plots, a black dot represents a participant, the red dashed line is the linear fit fitted using linear regression between the two variables and the grey patch represents the 95% confidence interval.

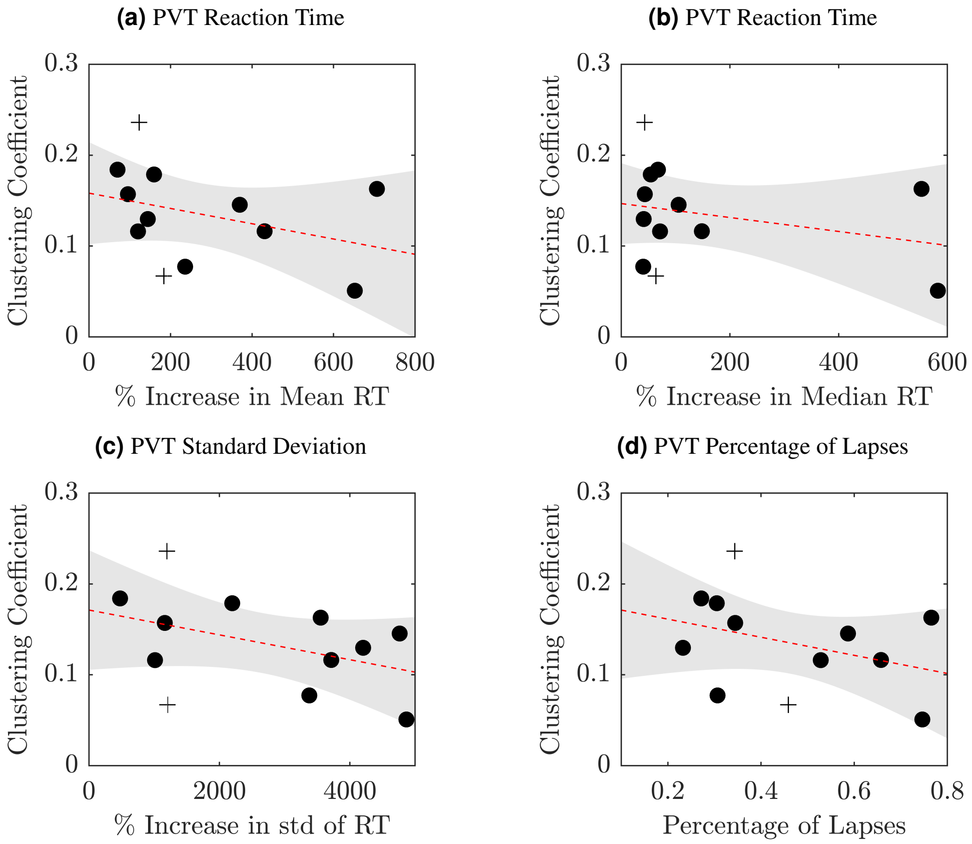


**Figure S7. Scatterplot Relating PVT Impairment to the Clustering Coefficient for the Acquisition 11 Hours Before DLMO.** Scatterplots relating the PVT impairment as measured using **(a)** mean RT, **(b)** median RT, **(c)** standard deviation of RT and **(d)** percentage of lapses to the clustering coefficient for the acquisition 11 hours before DLMO. For all plots, participants whose EEG acquisition 11 hours before DLMO is not their first recording are represented by +, while those whose first EEG acquisition is 11 hours before DLMO are denoted by • a black dot, the red dashed line is the linear fit fitted using linear regression between the two variables and the grey patch represents the 95% confidence interval.

#### Characteristic Path Length

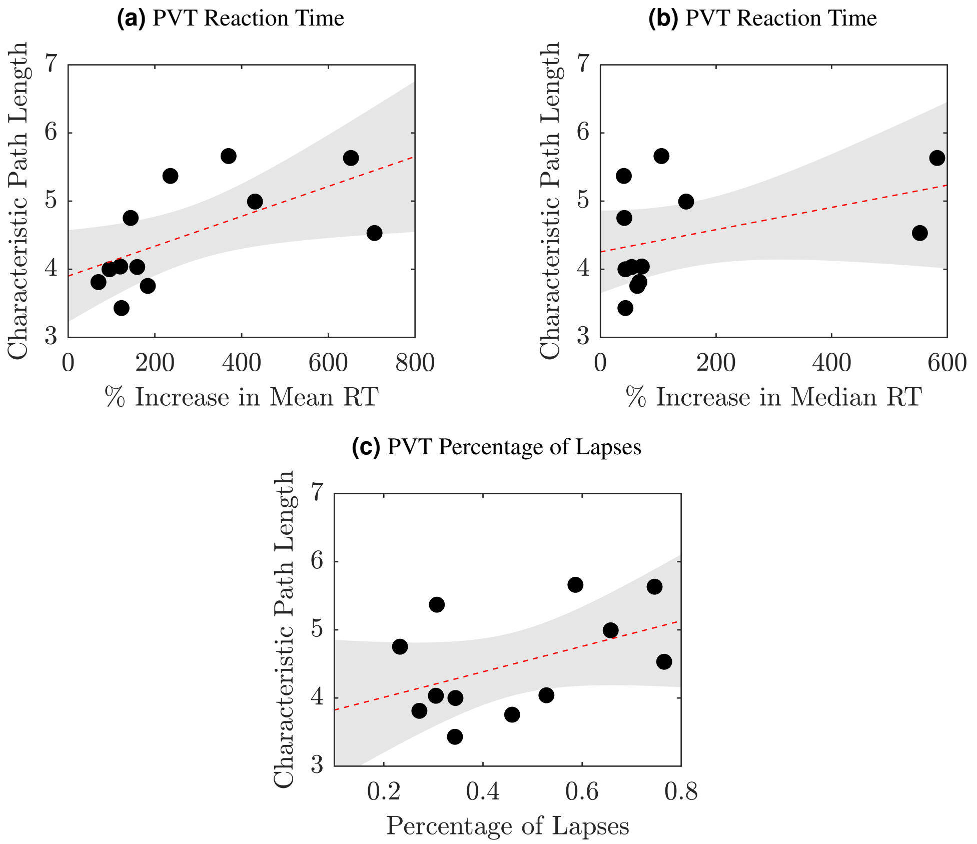


**Figure S8. Scatterplot Relating PVT Impairment to the Characteristic Path Length for the First Acquisition After Awakening.** Scatterplots relating the PVT impairment as measured using **(a)** mean RT, **(b)** median RT and **(c)** percentage of lapses to the characteristic path length for the first acquisition after awakening. For all plots, a black dot represents a participant, the red dashed line is the linear fit fitted using linear regression between the two variables and the grey patch represents the 95% confidence interval.

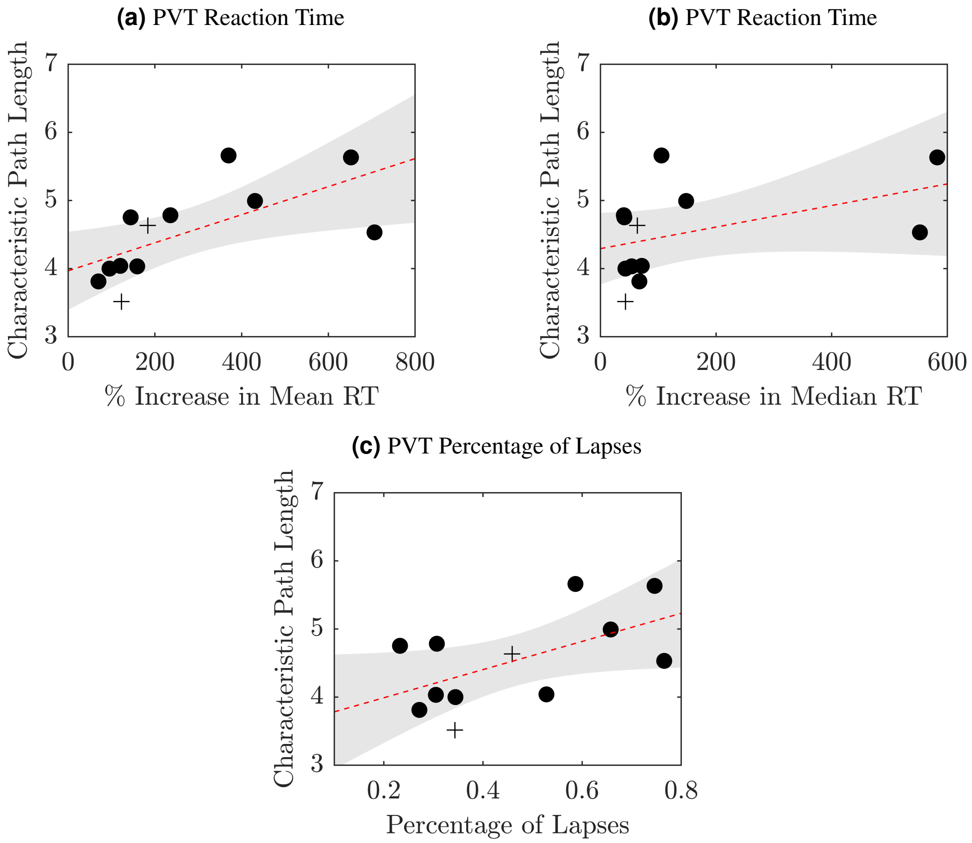


**Figure S9. Scatterplot Relating PVT Impairment to the Characteristic Path Length for the Acquisition 11 Hours Before DLMO.** Scatterplots relating the PVT impairment as measured using **(a)** mean RT, **(b)** median RT and **(c)** percentage of lapses to the characteristic path length for the acquisition 11 hours before DLMO. For all plots, participants whose EEG acquisition 11 hours before DLMO is not their first recording are represented by +, while those whose first EEG acquisition is 11 hours before DLMO are denoted by • a black dot, the red dashed line is the linear fit fitted using linear regression between the two variables and the grey patch represents the 95% confidence interval.

#### Stability

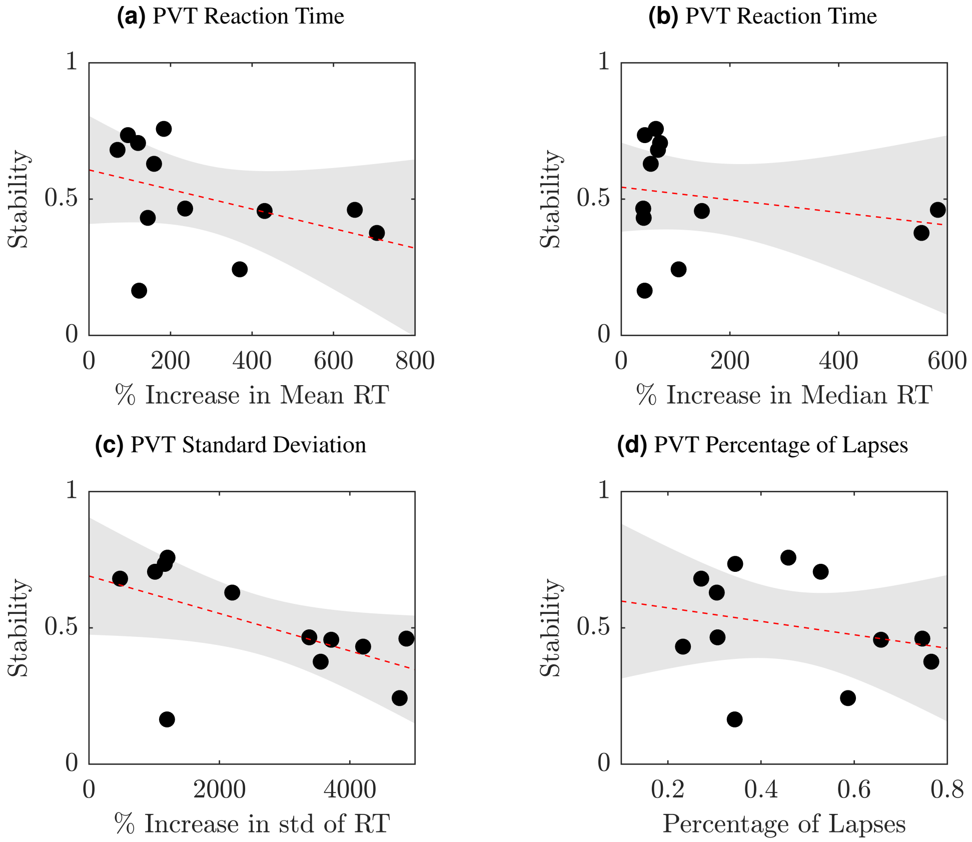


**Figure S10. Scatterplot Relating PVT Impairment to Stability for the First Acquisition After Awakening.** Scatterplots relating the PVT impairment as measured using **(a)** mean RT, **(b)** median RT, **(c)** standard deviation of RT and **(d)** percentage of lapses to stability for the first acquisition after awakening. For all plots, a black dot represents a participant, the red dashed line is the linear fit fitted using linear regression between the two variables and the grey patch represents the 95% confidence interval.

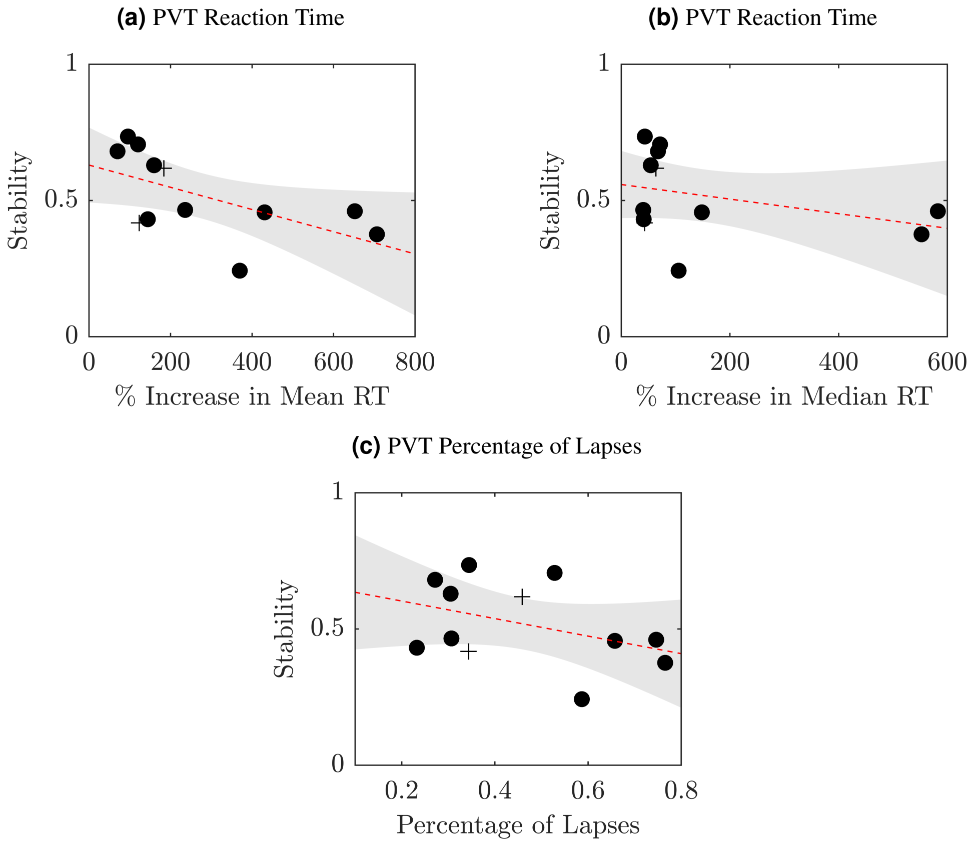


**Figure S11. Scatterplot Relating PVT Impairment to Stability for the Acquisition 11 Hours Before DLMO.** Scatterplots relating the PVT impairment as measured using **(a)** mean RT, **(b)** median RT and **(c)** percentage of lapses to stability for the acquisition 11 hours before DLMO. For all plots, participants whose EEG acquisition 11 hours before DLMO is not their first recording are represented by +, while those whose first EEG acquisition is 11 hours before DLMO are denoted by • a black dot, the red dashed line is the linear fit fitted using linear regression between the two variables and the grey patch represents the 95% confidence interval.

#### PVT Impairment: No Outliers

For the percentage increase in median RT two participants had an impairment measure more than 1.5 times the interquartile range away from the 75$^{th}$ percentile and were therefore considered outliers. The anlaysis was repeated with these participants removed and the Pearson's correlation and associated $p$-values are given in Supplementary Table S3.

| Baseline Acquisition: Graph Metric: PVT Impairment | Correlation | $p$-value | Adjusted $p$-value |
|---|---|---|---|
| $B_{FA}$: $\langle s \rangle$: Mean RT | -0.6784 | 0.0310 | 0.1116 |
| $B_{FA}$: $\langle s \rangle$: Median RT | -0.3051 | 0.3914 | 0.5219 |
| $B_{FA}$: $\langle s \rangle$: std$_{RT}$ | -0.8524 | 0.0017 | 0.0185* |
| $B_{FA}$: $\langle s \rangle$: % Lapses | -0.2500 | 0.4860 | 0.5760 |
| $B_{FA}$: $\langle C \rangle$: Mean RT | -0.4195 | 0.2276 | 0.4283 |
| $B_{FA}$: $\langle C \rangle$: Median RT | -0.1983 | 0.5829 | 0.6432 |
| $B_{FA}$: $\langle C \rangle$: std$_{RT}$ | -0.5252 | 0.1190 | 0.2798 |
| $B_{FA}$: $\langle C \rangle$: % Lapses | -0.2177 | 0.5458 | 0.6238 |
| $B_{FA}$ :$\lambda_L$: Mean RT | 0.7552 | 0.0116 | 0.0597 |
| $B_{FA}$ :$\lambda_L$: Median RT | 0.4244 | 0.2216 | 0.4283 |
| $B_{FA}$ :$\lambda_L$: std$_{RT}$ | 0.9006 | 0.0004 | 0.0121* |
| $B_{FA}$ :$\lambda_L$: % Lapses | 0.3477 | 0.3249 | 0.4950 |
| $B_{FA}$: $r_s$: Mean RT | -0.4345 | 0.2096 | 0.4283 |
| $B_{FA}$: $r_s$: Median RT | -0.1548 | 0.6693 | 0.6909 |
| $B_{FA}$: $r_s$: std$_{RT}$ | -0.5700 | 0.0853 | 0.2483 |
| $B_{FA}$: $r_s$: % Lapses | -0.1225 | 0.7360 | 0.7360 |
| $B_{-11}$: $\langle s \rangle$: Mean RT | -0.6681 | 0.0347 | 0.1116 |
| $B_{-11}$: $\langle s \rangle$: Median RT | -0.3601 | 0.3067 | 0.4950 |
| $B_{-11}$: $\langle s \rangle$: std$_{RT}$ | -0.7469 | 0.0131 | 0.0597 |
| $B_{-11}$: $\langle s \rangle$: % Lapses | -0.3515 | 0.3193 | 0.4950 |
| $B_{-11}$: $\langle C \rangle$: Mean RT | -0.3570 | 0.3112 | 0.4950 |
| $B_{-11}$: $\langle C \rangle$: Median RT | -0.1787 | 0.6214 | 0.6628 |
| $B_{-11}$: $\langle C \rangle$: std$_{RT}$ | -0.2910 | 0.4146 | 0.5307 |
| $B_{-11}$: $\langle C \rangle$: % Lapses | -0.3098 | 0.3837 | 0.5219 |
| $B_{-11}$ :$\lambda_L$: Mean RT | 0.8194 | 0.0037 | 0.0296* |
| $B_{-11}$ :$\lambda_L$: Median RT | 0.5428 | 0.1050 | 0.2798 |
| $B_{-11}$ :$\lambda_L$: std$_{RT}$ | 0.8569 | 0.0015 | 0.0185* |
| $B_{-11}$ :$\lambda_L$: % Lapses | 0.5212 | 0.1224 | 0.2798 |
| $B_{-11}$: $r_s$: Mean RT | -0.6677 | 0.0349 | 0.1116 |
| $B_{-11}$: $r_s$: Median RT | -0.3183 | 0.3701 | 0.5219 |
| $B_{-11}$: $r_s$: std$_{RT}$ | -0.8086 | 0.0046 | 0.0296* |
| $B_{-11}$: $r_s$: % Lapses | -0.2715 | 0.4480 | 0.5513 |

**Table S3. Results of correlating the Baseline Graph Metric with PVT Impairment When Outliers Were Removed.** The Pearson's correlation between the participants' PVT impairment and the graph metric for both baseline acquisitions when correlating mean node strength ($\langle s \rangle$), clustering coefficient ($\langle C \rangle$), characteristic path length ($\lambda_L$) and stability ($r_s$) with the four PVT measures after the two outliers were removed. The $p$-values were corrected for multiple comparisons using the Benjamini-Hochberg correction for FDR[1]. * denotes significance $p$-value < 0.05.

### PVT Performance

#### Mean Node Strength

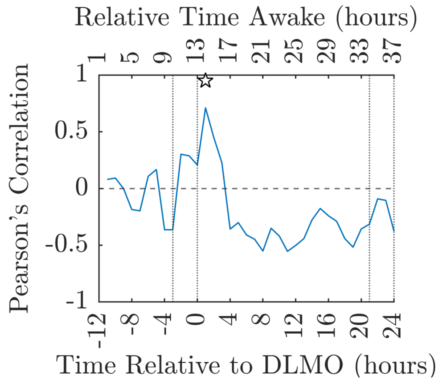


**(a)** Mean PVT RT (ms)

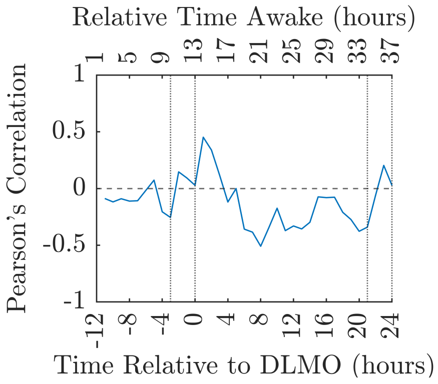


**(b)** Median PVT RT (ms)

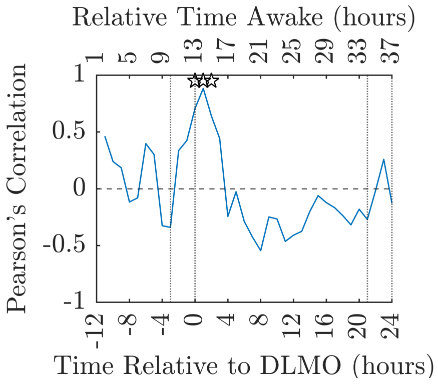


**(c)** PVT Percentage of Lapses

**Figure S12. Pearson's Correlation Between Baseline Mean Node Strength for the First Acquisition and PVT Performance.** How the Pearson's correlation coefficient between the baseline mean node strength for the first acquisition after awakening and the PVT performance as measured using **(a)** mean RT, **(b)** median RT, **(c)** standard deviation of RT and **(d)** percentage of lapses changes over the CR. In all plots the ☆ indicates where the $p$-value associated with the correlation is significant $p < 0.05$ (uncorrected) and the dotted vertical lines indicate the times considered to be in the WMZ (3 hours before DLMO to 5 minutes after) and the relative time awake provides an indication of the time awake for the majority of the participants.

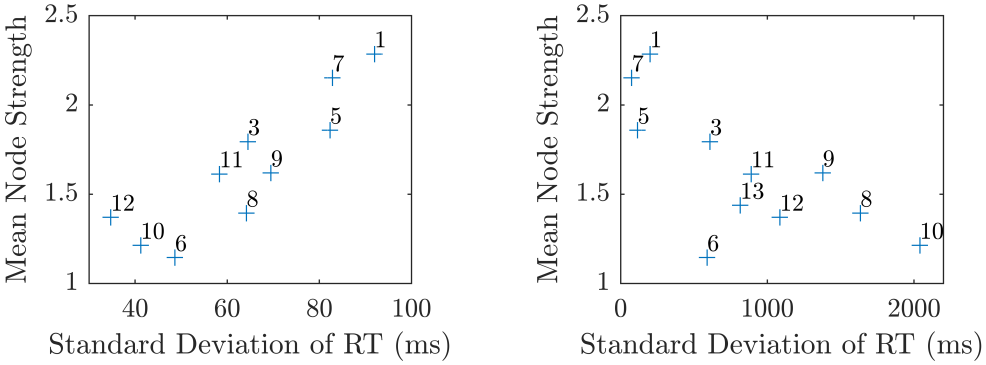


**(a)** DLMO + 1 h

**(b)** DLMO + 11 h

**Figure S13. Scatterplots Relating Mean Node Strength to Standard Deviation of RT for the First Acquisition.** The relationship between mean node strength and standard deviation of RT at **(a)** DLMO + 1 h and **(b)** DLMO + 11 h where participants are individually labelled. Note Participant 4 had data missing for these acquisitions.

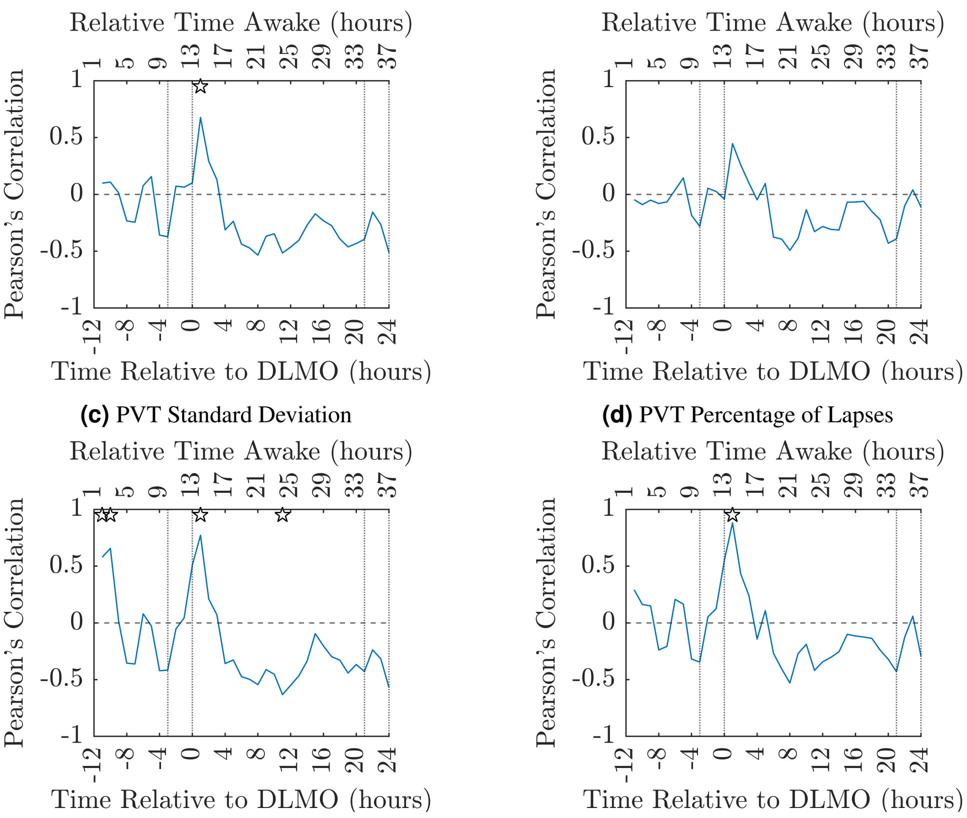


**(a)** Mean PVT RT (ms)

**(b)** Median PVT RT (ms)

**(c)** PVT Standard Deviation

**(d)** PVT Percentage of Lapses

**Figure S14. Pearson's Correlation Between Baseline Mean Node Strength for the Acquisition 11 hours before DLMO and PVT Performance.** How the Pearson's correlation coefficient between the baseline mean node strength for the acquisition 11 hours before DLMO and the PVT performance as measured using **(a)** mean RT, **(b)**, median RT **(c)** standard deviation of RT and **(d)** percentage of lapses changes over the CR. In all plots the ☆ indicates where the $p$-value associated with the correlation is significant $p < 0.05$ (uncorrected) and the dotted vertical lines indicate the times considered to be in the WMZ (3 hours before DLMO to 5 minutes after) and the relative time awake provides an indication of the time awake for the majority of the participants.

#### Clustering Coefficient

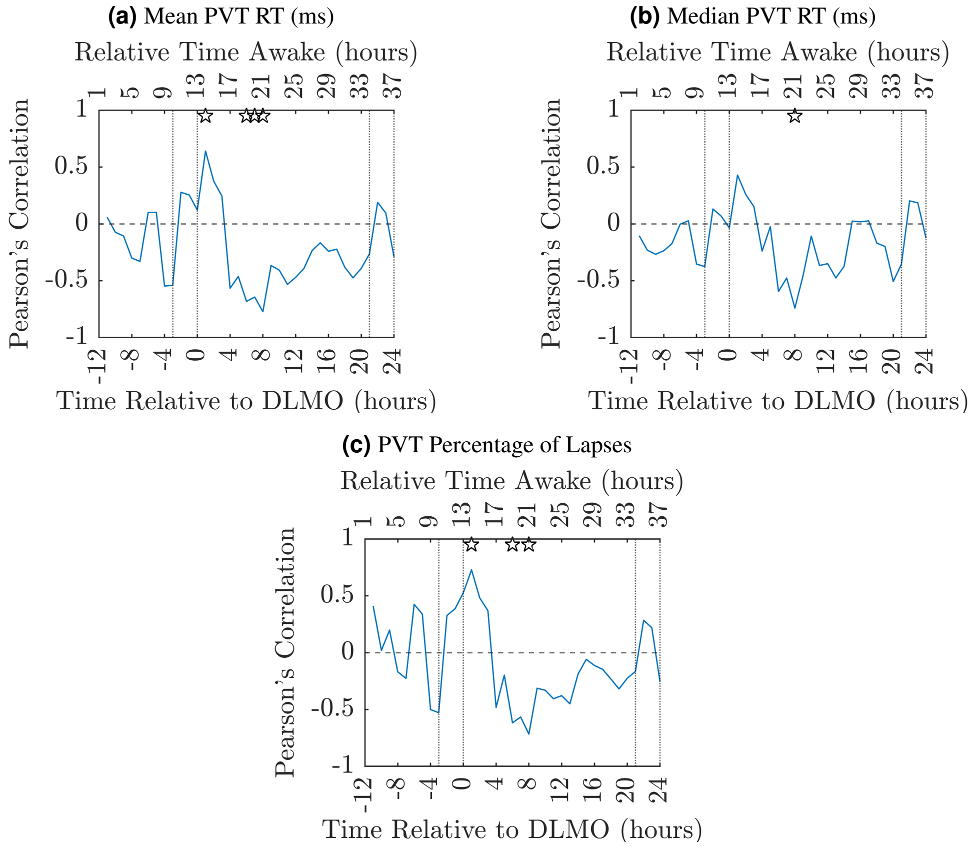


**Figure S15. Pearson's Correlation Between Baseline Clustering Coefficient for the First Acquisition and PVT Performance.** How the Pearson's correlation coefficient between the baseline clustering coefficient for the first acquisition after awakening and the PVT performance as measured using **(a)** mean RT, **(b)** median RT, **(c)** standard deviation of RT and **(d)** percentage of lapses changes over the CR. In all plots the ☆ indicates where the *p*-value associated with the correlation is significant $p < 0.05$ (uncorrected) and the dotted vertical lines indicate the times considered to be in the WMZ (3 hours before DLMO to 5 minutes after) and the relative time awake provides an indication of the time awake for the majority of the participants.

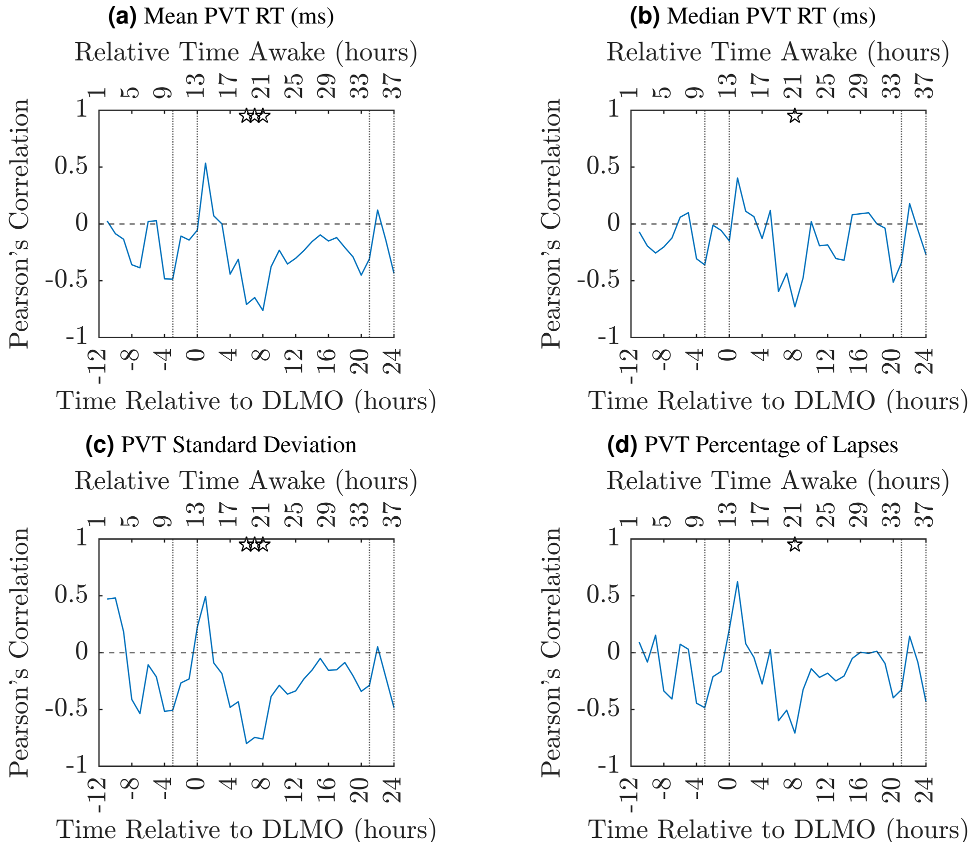


**Figure S16.** **Pearson's Correlation Between Baseline Clustering Coefficient for the Acquisition 11 hours before DLMO and PVT Performance.** How the Pearson's correlation coefficient between the baseline clustering coefficient for the acquisition 11 h before DLMO and the PVT performance as measured using **(a)** mean RT, **(b)** median RT, **(c)** standard deviation of RT and **(d)** percentage of lapses changes over the CR. In all plots the ☆ indicates where the *p*-value associated with the correlation is significant $p < 0.05$ (uncorrected) and the dotted vertical lines indicate the times considered to be in the WMZ (3 hours before DLMO to 5 minutes after) and the relative time awake provides an indication of the time awake for the majority of the participants.

#### Characteristic Path Length

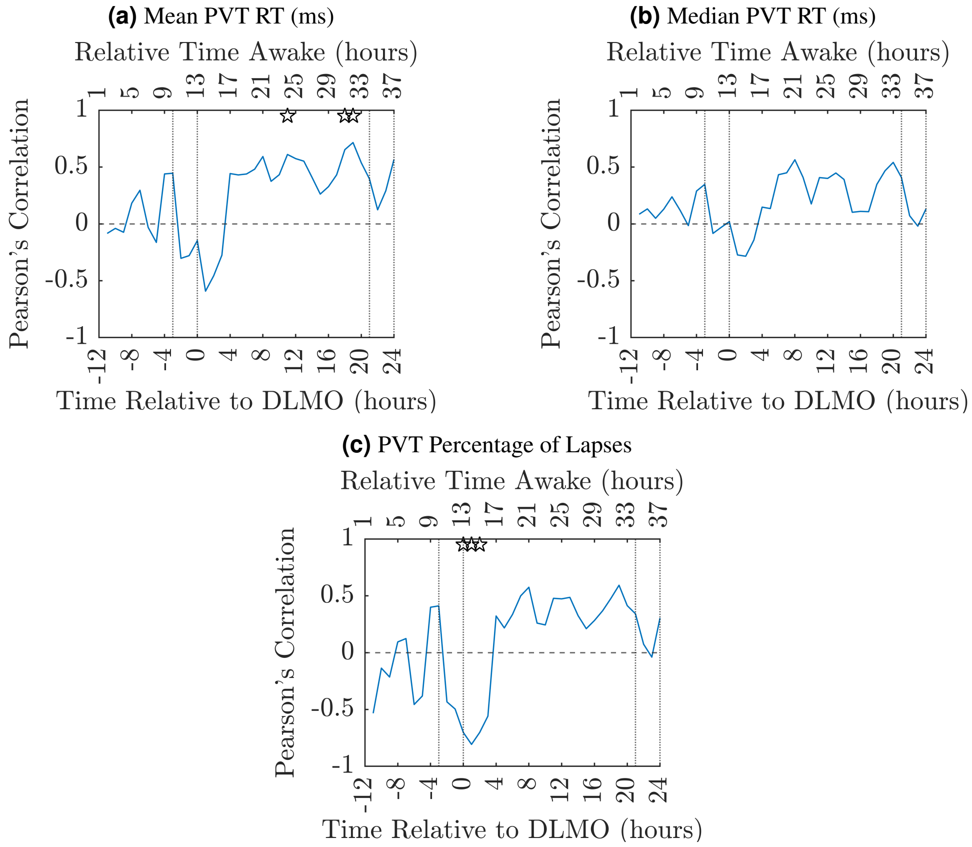


**Figure S17. Pearson's Correlation Between Baseline Characteristic Path Length for the First Acquisition and PVT Performance.** How the Pearson's correlation coefficient between the baseline characteristic path length for the first acquisition after awakening and the PVT performance as measured using **(a)** mean RT, **(b)** median RT, **(c)** standard deviation of RT and **(d)** percentage of lapses changes over the CR. In all plots the ☆ indicates where the *p*-value associated with the correlation is significant $p < 0.05$ (uncorrected) and the dotted vertical lines indicate the times considered to be in the WMZ (3 hours before DLMO to 5 minutes after) and the relative time awake provides an indication of the time awake for the majority of the participants.

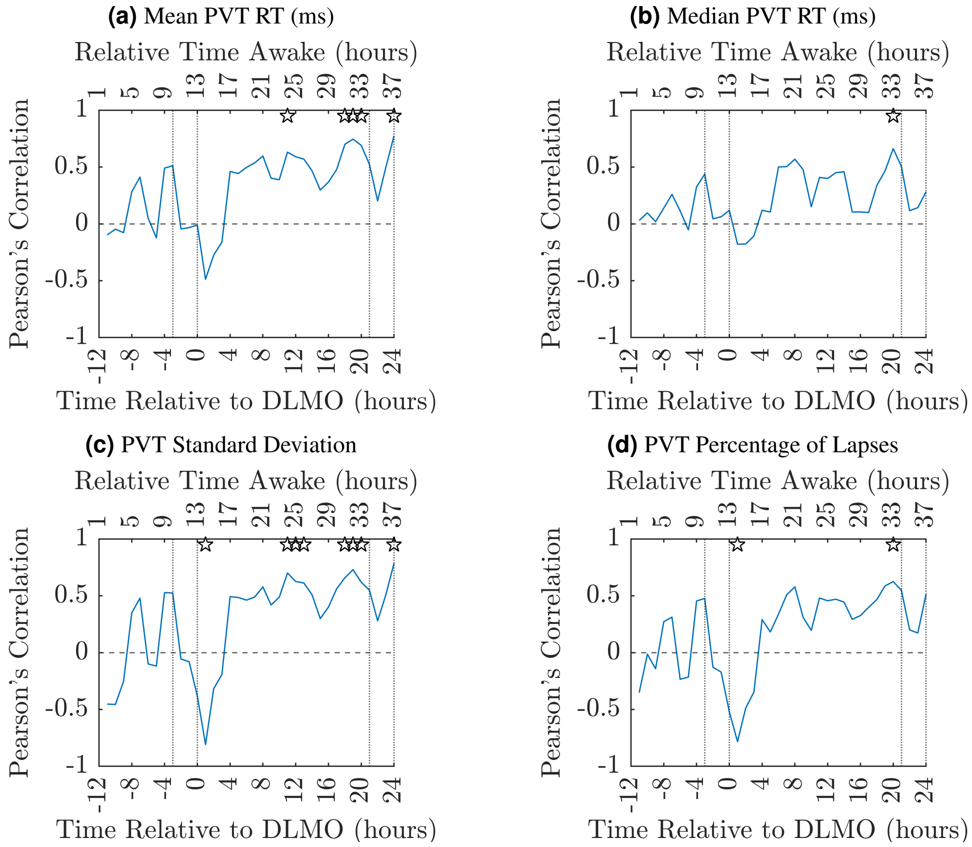


**Figure S18. Pearson's Correlation Between Baseline Characteristic Path Length for the Acquisition 11 hours before DLMO and PVT Performance.** How the Pearson's characteristic path length between the baseline clustering coefficient for the acquisition 11 h before DLMO and the PVT performance as measured using **(a)** mean RT, **(b)** median RT, **(c)** standard deviation of RT and **(d)** percentage of lapses changes over the CR. In all plots the ☆ indicates where the $p$-value associated with the correlation is significant $p < 0.05$ (uncorrected) and the dotted vertical lines indicate the times considered to be in the WMZ (3 hours before DLMO to 5 minutes after) and the relative time awake provides an indication of the time awake for the majority of the participants.

#### Stability

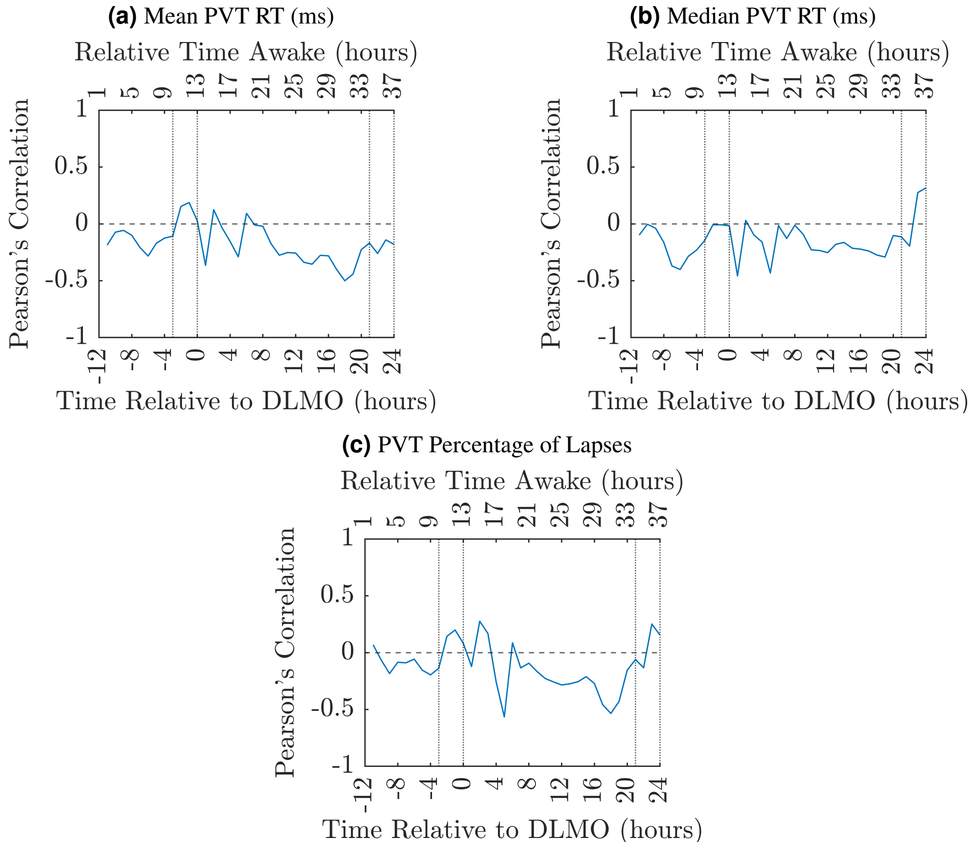


**Figure S19.** **Pearson's Correlation Between Baseline Stability for the First Acquisition and PVT Performance.** How the Pearson's correlation coefficient between the baseline stability for the first acquisition after awakening and the PVT performance as measured using **(a)** mean RT, **(b)** median RT, **(c)** standard deviation of RT and **(d)** percentage of lapses changes over the CR. In all plots the ☆ indicates where the $p$-value associated with the correlation is significant $p < 0.05$ (uncorrected) and the dotted vertical lines indicate the times considered to be in the WMZ (3 hours before DLMO to 5 minutes after) and the relative time awake provides an indication of the time awake for the majority of the participants.

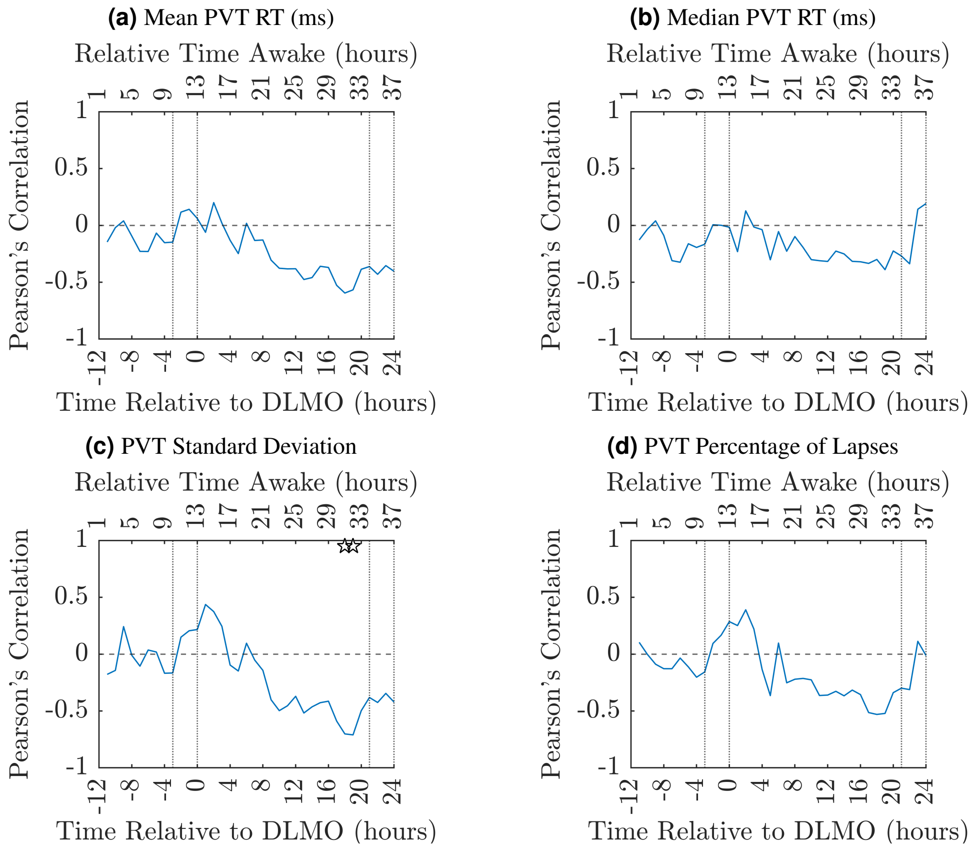


**Figure S20. Pearson's Correlation Between Baseline Stability for the Acquisition 11 hours before DLMO and PVT Performance.** How the Pearson's correlation coefficient between the baseline stability for the acquisition 11 h before DLMO and the PVT performance as measured using **(a)** mean RT, **(b)** median RT, **(c)** standard deviation of RT and **(d)** percentage of lapses changes over the CR. In all plots the ☆ indicates where the $p$-value associated with the correlation is significant $p < 0.05$ (uncorrected) and the dotted vertical lines indicate the times considered to be in the WMZ (3 hours before DLMO to 5 minutes after) and the relative time awake provides an indication of the time awake for the majority of the participants.

#### Stability

##### Group Level

| Frequency Band | Paired Cohen's $d$ | $p$-value | Adjusted $p$-value |
|---|---|---|---|
| Delta | 0.0952 | 0.9495 | 0.9495 |
| Theta | 0.2115 | 0.0989 | 0.3455 |
| Alpha | 0.2326 | 0.1727 | 0.3455 |
| Beta | 0.1124 | 0.5592 | 0.7455 |

**Table S4. Stability Prior to and During the WMZ.** Cohen's $d$ effect size and associated $p$-values for paired $t$-tests for each frequency band. The tests compared the mean stability for 3 hours prior to the WMZ to the WMZ with the mean stability for the participant over the 3 hours during the WMZ. Both $p$-values and adjusted $p$-values corrected for multiple comparisons using the Benjamini-Hochberg method are given[1].

##### PVT Groups

| Frequency Band | DLMO-11h - DLMO | DLMO+1h - DLMO+12h | DLMO+13h - DLMO+24h |
|---|---|---|---|
| Delta | 0.1136 (0.0947) | 0.1179 (0.1081) | 0.4937 (0.4937) |
| Theta | $0.0010^{*}$ (0.0004) | 0.1132 (0.0849) | $0.0010^{*}$ (0.0004) |
| Alpha | $< 0.0001^{*}$ ($< 0.0001$) | $0.0030^{*}$ (0.0015) | $< 0.0001^{*}$ ($< 0.0001$) |
| Beta | $< 0.0001^{*}$ ($< 0.0001$) | 0.0694 (0.0405) | 0.0739 (0.0493) |

**Table S5. Table Showing the $p$-values for Permutation Tests Comparing Stability with Participants Grouped by Standard Deviation of RT.** Results for permutation tests ($n = 10,000$) comparing the mean value of the low and high impairment groups over adjacent 12 hour periods. The $p$-values have been corrected using the Benjamini-Hochberg method of correcting for FDR[1]. $^{*}$ denotes significance $p$-value $< 0.05$. Uncorrected $p$-values are given in brackets.

#### Individual

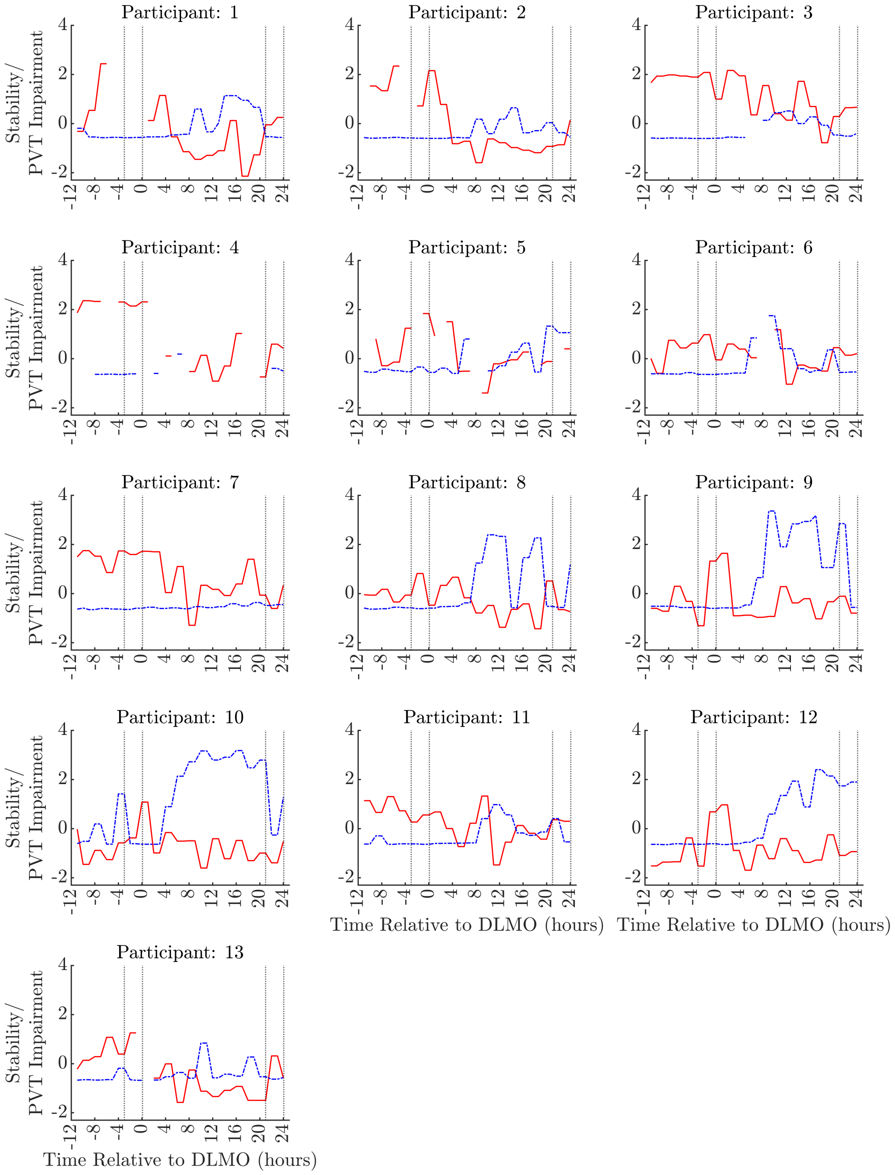


**Figure S21. Individual Stability and Standard Deviation on PVT.** The individual time-series for the standard deviation of RT on the PVT (-·-·-·-) and the stability of the FNs (——). The dotted vertical lines indicate the times considered to be in the WMZ (3 hours before DLMO to 5 minutes after).

| Participant | Impairment Group | Spearman's Rank Correlation | $p$-value | Adjusted $p$-value |
|---|---|---|---|---|
| 1 | Low | -0.7571 | < 0.0001 | < 0.0001* |
| 2 | Low | -0.7527 | < 0.0001 | < 0.0001* |
| 3 | Low | -0.6603 | < 0.0001 | < 0.0001* |
| 4 | Low | -0.8261 | 0.0032 | 0.0052* |
| 5 | Low | -0.1862 | 0.3334 | 0.3940 |
| 6 | High | -0.3015 | 0.0831 | 0.1200 |
| 7 | Low | -0.5418 | 0.0006 | 0.0017* |
| 8 | High | -0.5055 | 0.0017 | 0.0036* |
| 9 | High | -0.1977 | 0.2478 | 0.3222 |
| 10 | High | -0.1363 | 0.4279 | 0.4318 |
| 11 | NA | -0.4923 | 0.0023 | 0.0043* |
| 12 | High | 0.1352 | 0.4318 | 0.4318 |
| 13 | High | -0.5602 | 0.0006 | 0.0017* |

**Table S6. Spearman's Rank Correlation Between std RT and Stability for Individuals.** The Spearman's rank correlation and associated $p$-value for the correlation between the participants standard deviation of RT and the stability of their FNs over the CR. The $p$-values have been corrected using the Benjamini-Hochberg method of correcting for FDR[1]. * denotes significance $p$-value < 0.05.

#### Stability: Additional Participants

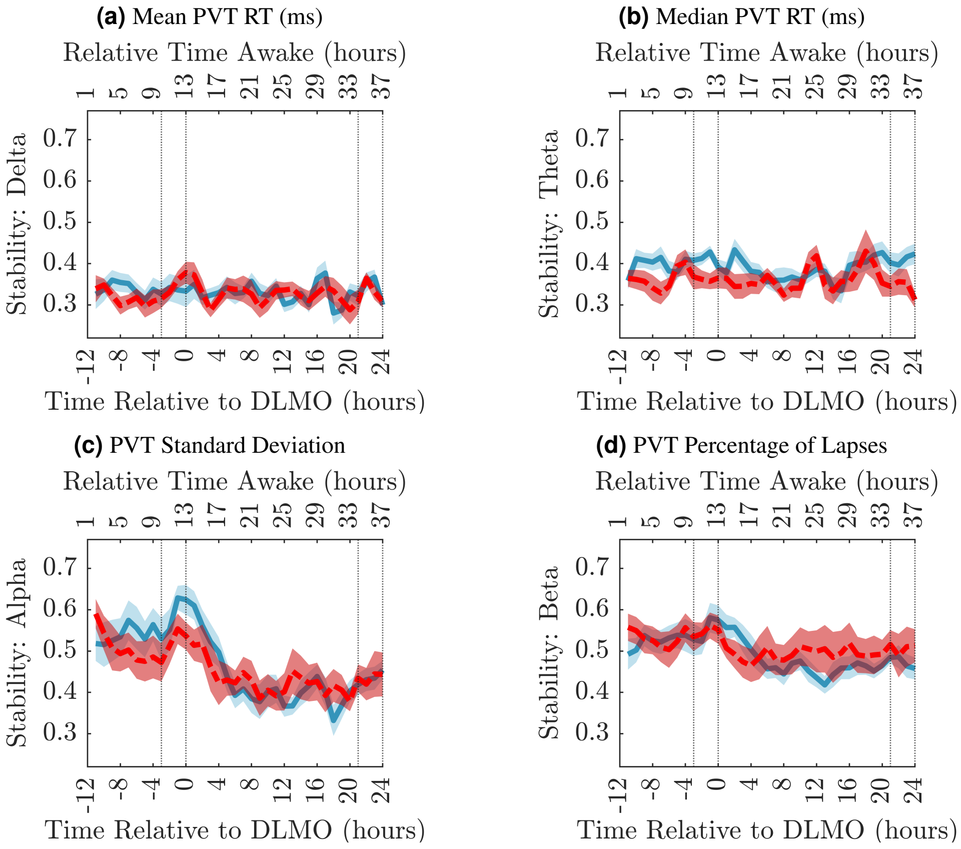


**Figure S22. Stability of Participants FN's: Additional Participants.** The median stability, as measured from correlating the PLF networks from all epochs for that participant from an EEG acquisition for a given frequency band **(a)** Delta, **(b)** Theta, **(c)** Alpha and **(d)** Beta. The solid blue line (——) is the mean across the median stability of the 13 participants in this study and the dashed red line (- - - - -) is the mean across the median stability of the 9 additional participants. The corresponding blue and red shaded area is the standard error of the mean. The dotted vertical lines indicate the times considered to be in the WMZ (3 hours before DLMO to 5 minutes after).
